# The Deubiquitinating Enzyme Otub2 Modulates Pancreatic Beta-Cells Function and Survival

**DOI:** 10.1101/2024.09.30.615641

**Authors:** Michal Oshry, Roi Isaac, Sigalit Boura-Halfon, Sanford Sampson, Sima Lev, Yehiel Zick, Yaron Vinik

## Abstract

We have previously shown that otubain 2 (OTUB2), a deubiquitinating enzyme, inhibits caspase-3/7 activity in primary human islets; promotes insulin secretion and inhibits cytokine-induced nuclear factor-κB (NFκB) activity. In the present work we show that overexpression of Otub2 in MIN6 cells inhibits NFκB activity and the expression of its target genes MCP-1 and iNOS. Consequently, both the basal and the cytokine-induced apoptosis of cultured MIN6 cells and dispersed human islets were inhibited. Overexpression of Otub2 in MIN6 cells increase the mRNA levels of NKx6.1 and Glut2 and concomitantly increased glucose-stimulated insulin secretion (GSIS) (by 2-3-fold). The beneficial effects of Otub2 on β-cell function was demonstrated by the phenotype of *Otub2*^-/+^ and *Otub2*^-/-^ mice, which manifested impaired glucose tolerance and increased expression of NFkB target genes (e.g. IP-10, MCP-1 and IL-1β). RNAseq analysis of pancreata derived from OTUB2 KO mice revealed reduced expression of genes that down regulate K^+^ transporters (e.g. *Ank2*, *Cacna1a* and *Kcnab1)* combined with an increase in oxidative phosphorylation related genes. Given that closure of K^+^ channels is crucial for insulin secretion, these results could account, at least in part, for the impaired GSIS in the OTUB2 KO mice. Indeed, mass-spectrometry analysis of proteins co-immunoprecipitated with Otub2 revealed the voltage-gated potassium channel subunit Kv9.3 as a major Otub2 binding-partner. Additional binding partners included the Peg3 and Camk2d proteins, which promote NFκB signaling and β-cell death. Hence, by deubiquitinating proteins in complexes that contain Peg3 and Camk2d, Otub2 might inhibit propagation of NFκB signaling and β-cell apoptosis. Collectively our findings implicate Otub2 as a key regulator of β cell function, mainly affecting NFkB signaling and the K^+^ channels that regulate insulin secretion.

## Introduction

Type-1 and Type-2 diabetes mellitus involve selective and progressive loss of pancreatic β-cells; thus, both forms of the disease might eventually require life-long insulin replacement therapy [1, 2]. Transplantation of islets and β-cells regeneration are two major approaches for β-cells replenishment [3]. Currently, islets transplantation is the preferred treatment that brings about insulin-independence, improves quality of life and increased beta-score [4–6]. However, a large proportion (up to 70%) of the graft is lost during the initial few days following islet infusion. This is mainly attributed to inflammatory reactions, reactive oxygen species generation, and cytokine secretion, all challenges that islets face following transplantation [5, 7]. For example, oxygen tension in liver, the preferred site for islet transplantation, is well below that of the pancreas. Further, the low expression level of major cellular antioxidant enzymes in islets, contributes to less-than-optimal islet survival. Autoimmune and alloimmune responses and the diabetogenic action of immunosuppressive drugs also contribute to islet deterioration [5]. Several strategies have been employed to improve graft survival, such as heparinization of the islets’ medium prior to transplantation and peri-transplant insulin therapy. These approaches may improve beta-cell function and reduce immediate beta-cell apoptosis, but cannot prevent loss of beta cell function over time, with most patients returning to insulin-dependence after five-years [4, 8].

To improve beta-cells survival following transplantation, we have previously developed and performed high throughput screens (HTS) of ∼730 pre-selected siRNA in search for genes that affect survival of isolated human pancreatic islets treated with pro-inflammatory cytokines [9, 10]. These studies identified a number of novel genes (e.g. Otub2 [9], Ndfip1 [11] and TM7SF3 [12], whose roles in survival of cytokine-treated human β-cells had not been previously examined.

Ovarian tumor (Otu) domain-containing ubiquitin aldehyde-binding protein (Otub2), a family member of cysteine proteases having a deubiquitinase activity [13], was a highly significant ‘hit’ in the above screens. Silencing of Otub2 expression increased caspase-3/7 activity in primary human islets treated with a mixture of cytokines (TNFα, IL-1β IFNγ), inhibited insulin secretion, and increased NFκB activity [9]. These findings suggest that Otub2 may function as a novel protector of viability and insulin secretion in β-cells treated with cytokines. Otub2, as opposed to its close homologue Otub1, was shown to have a clear preference for cleaving K63–linked ubiquitin [14]. In doing so, Otub2 could hamper NFκB activation by modifying scaffold elements. In this study, we demonstrate the beneficial effects of Otub2 *in-vivo* and show its action as a pro-survival protein for β-cells that protects them from apoptotic death; preserves their mass and maintains their functionality. These findings suggest that Otub2 might be a potential candidate for improved survival of transplanted beta-cell.

## Materials and Methods

### Cells

MIN6 murine pancreatic β-cell line (passages 25-39) were cultured in Dulbecco’s modified Eagle’s medium (DMEM) containing 11 mM glucose supplemented with 10% FBS, 2 mM L-glutamine and 5 μM β-mercaptoethanol. 100 U/ml penicillin and 100 μg/ml streptomycin were added to non-transfected MIN6 cells, while 400 μg/ml geneticin (G418) was added to stably transfected MIN6 cells. All cells were grown at 37°C in a 5% CO_2_ humidified atmosphere. Mycoplasma was monitored pariodically.

### Plasmids

pEGFP-c1 plasmid (Addgene) harboring G418 resistance was used for creating stable MIN6 cell lines that overexpress Otub2 (mouse isoform 2) with Flag tag (instead of EGFP). pEGFP-c1 vector in which Flag tag was cloned instead of EGFP sequence (pFlag) served as control. pEGFP-c1 plasmids which contained the original EGFP sequence (p-EGFP-c1) and EGFP-Otub2 sequence (pEGFP-Otub2) were used for transient transfection of MIN6 cells. To create these constructs, the Otub2 cDNA was amplified by PCR using the 5’ primer (5’-CAGTCCGGAGACACTATGAGTGAAACATCTTTCAACC-3’); and the 3’ primer (5’-CGCGGATCCGCGGTAGTCAGTGTTTCTCGGCTGC-3’). The PCR product was digested using BspE1 and BamHI restriction enzymes and ligated into pEGFP-c1 plasmid. An additional construct was generated using the following primers containing Flag tag: 5’ primer (5’-CTAGCATGGACTACAAAGACGATGACGACAAGT-3’) and 3’ primer (5’-CCGGACTTGTCGTCATCGTCTTTGTAGTCCATG-3’) the primers were hybridized and ligated into pEGFP-c1 digested with NheI and BspEI restriction enzymes. The EGFP cDNA was then excised from the plasmid and replaced by Flag tag creating pFlag-Otub2 construct. pFlag plasmid was generated after digestion of pFlag-Otub2 construct with BspE1 and BamHI restriction enzymes (excising Otub2 out of the construct) and used as control.

### Antibodies

Polyclonal anti-Flag antibodies were from Sigma. Polyclonal anti-Otub2 antibodies were from Novus Biologicals and Abcam. Polyclonal IKKβ phosphorylated antibodies were from Cell Signaling Technology. Polyclonal anti-GFP antibodies were from Abcam. Polyclonal anti-GAPDH antibodies were from EMD Millipore. Alexa488 conjugated goat anti-mouse secondary antibodies were from Life Technologies.

### Cytokines

The cytokine mixture referred to as ‘1x-cytomix’ consisted of 3nM TNF-α; 3nM γ-IFN and 1.5nM IL1-β. Their biological activities were 10 units/ng (TNFα, IFNγ) and 200 units/ng (IL-1β).

### Dispersion and culture of human islets

Isolated human islets (∼90% purity confirmed by dithizone staining) were provided by the European Consortium for Islets Transplantation (Islet for Basic Research program) through a Juvenile Diabetes Research Foundation Award 31-2008-413. Islets were cultured at 37°C in a 5% CO2 humidified atmosphere in CMRL 1066 medium containing 10% (v/v) FBS, 2mM L-glutamine, 100 units/ml penicillin, 100 μg/ml streptomycin, 0.25 μg/ml amphotericin, and 40 μg/ml gentamycin. The medium was changed every other day. Intact human islets were dispersed by four minutes incubation at 37°C with 1mg/ml Trypsin/EDTA by pipetting the cells, and by passing them twice, through a 21G needle. Trypsinized islets were washed with CMRL 1066 medium containing 10% FBS and were resuspended in CMRL 1066 containing 10% FBS. Cells were used within 48 hours following dispersion. Human islets studies received Ethics Committee approval.

### siRNA transfection

MIN6 cells or human islets were seeded in 96 well plates (30,000 cells/well or 1000 islets/well) in 100µl medium (DMEM for MIN6 cells and CMRL for human islets), and immediately transfected with sequences of non-targeting siRNA or Otub2 siRNA (to a final concentration of 25nM) using DarmaFECT-4 transfection reagent for MIN6 cells and Darma-FECT-1 transfection reagent for human islets, according to manufacturer’s instructions (Dharmacon).

### Transient DNA transfection

MIN6 cells or human islets were seeded in plates (500,000 cells/well in a 6-well plates for MIN6 cells, and 1000 islets/well in a 96-well plates for human islets). 24 hours post seeding, the cells or human islets were transfected with pFlag or pFlag-Otub2 vectors (3 µg/well for a 6-well plate and 0.2 µg/well for a 96-well plate) using Lipofectamine 2000 transfection reagent according to manufacturer’s instructions (Thermo-Fisher Scientific).

### Generation of MIN6 cells that stably over-express Otub2

MIN6 cells were seeded in 12-well plates. 24 hours post seeding cells were transfected, using jetPEI transfection reagent (Polyplus Transfection), with pEGFP-c1 vectors (1µg/well) harboring G418 resistance to calibrate for the cells’ G418 resistance. pEGFP-c1 vector in which Flag tag was cloned instead of EGFP sequence (pFlag) served as control; and pEGFP-c1 with Flag-Otub2 cDNA clone without EGFP sequence (pFlag-Otub2) was utilized to overexpress Otub2. Non-transfected cells served as control. 48 hours after transfection media were replaced with media containing elevated concentrations of G418. After the calibration process was complete, and the proper concentration of G418 required for selection was established, MIN6 cells were seeded in 6-well plates (150,000 cells/well) to create stable cell lines. 24 hours post-seeding, cells were transfected with pEGFP-c1 constructs (3 µg/well). 48 hours thereafter the media were replaced with media containing 400µg/ml G418 for selection process. After completion of selection, media were replaced with media containing 200µg/ml G418 for maintenance.

### Caspase 3/7 activity assay

MIN6 cells were seeded in 96-well plates and transfected with siRNA as described above. 48 hours after transfection, cells were treated with cytokine mixture, referred to as 1x-cytomix. For stably transfected cell lines, cells were treated with 1x-cytomix 24 hours post seeding. 24 hours later caspase 3/7 activity was measured using Sensolyte homogeneous RH110 caspase 3/7 assay kit (AnaSpec). Human islets were dispersed and seeded in 96-well plates (1000 islets/well). Islets were transfected with p-Flag or pFlag-Otub2 constructs that stably express Flag-Otub2. The rest of the experiment was similar to the assay in MIN6 cells as described above.

### NF-κB activity assay

NF-κB activity was determined using the Ready-To-Glow secreted luciferase assay kit (Clontech). Stably transfected MIN6 cells were seeded in 96-well plates (30,000 cells/well). 24 hours post seeding cells were transfected with a secreted luciferase plasmid (250 ng/well), coupled to the NF-κB enhancer elements. Constitutively secreted luciferase plasmid, transfected to separate wells (100ng/well), served as control. Twenty-four hours thereafter cells were treated with 1x-cytomix for 4 hours, to induce NF-κB activity. 50µl of the cells’ media were harvested and transferred to black 96-well plates containing a luciferase assay mix (5μl/well) added in advance. Luminescence was read using Infinite 2000 PRO multimode reader (Tecan Trading).

### Glucose stimulated insulin secretion (GSIS)

Stably transfected MIN6 cells were seeded in 96-well plates (30,000 cells/well). After 48 hours cells were incubated for 1 hour in Krebs-Ringer bicarbonate HEPES buffer (KRBH), containing 124mM NaCl, 5.6mM KCl, 2.5mM CaCl2 and 20mM HEPES pH 7.4, at 37°C, followed by incubation for 1 hour in KRBH with 0 mM or 20 mM glucose. Insulin concentration in the culture medium was determined using insulin detecting HTRF kit (Cisbio) according to manufacturer’s instructions.

### RNA analysis

Cells were grown in 24-well plates. After treatment, cells were harvested and total RNA was extracted using the PerfectPure RNA kit (5-prime, MD). RNA was quantified using nanodrop, and cDNA was generated by cDNA Reverse Transcription kit (Applied Biosystems, CA), following the manufacturer instructions. Quantitative detection of mRNA transcripts was carried out by real-time PCR using ABI-Prism 7300 instrument (Applied Biosystems, CA); SYBR Green PCR mix (Invitrogen) and specific primers (100 nM final concentration). Results were normalized to mRNA levels of Actin or HPRT. Primers used are given herein (see Table 1).

**Table 1.**
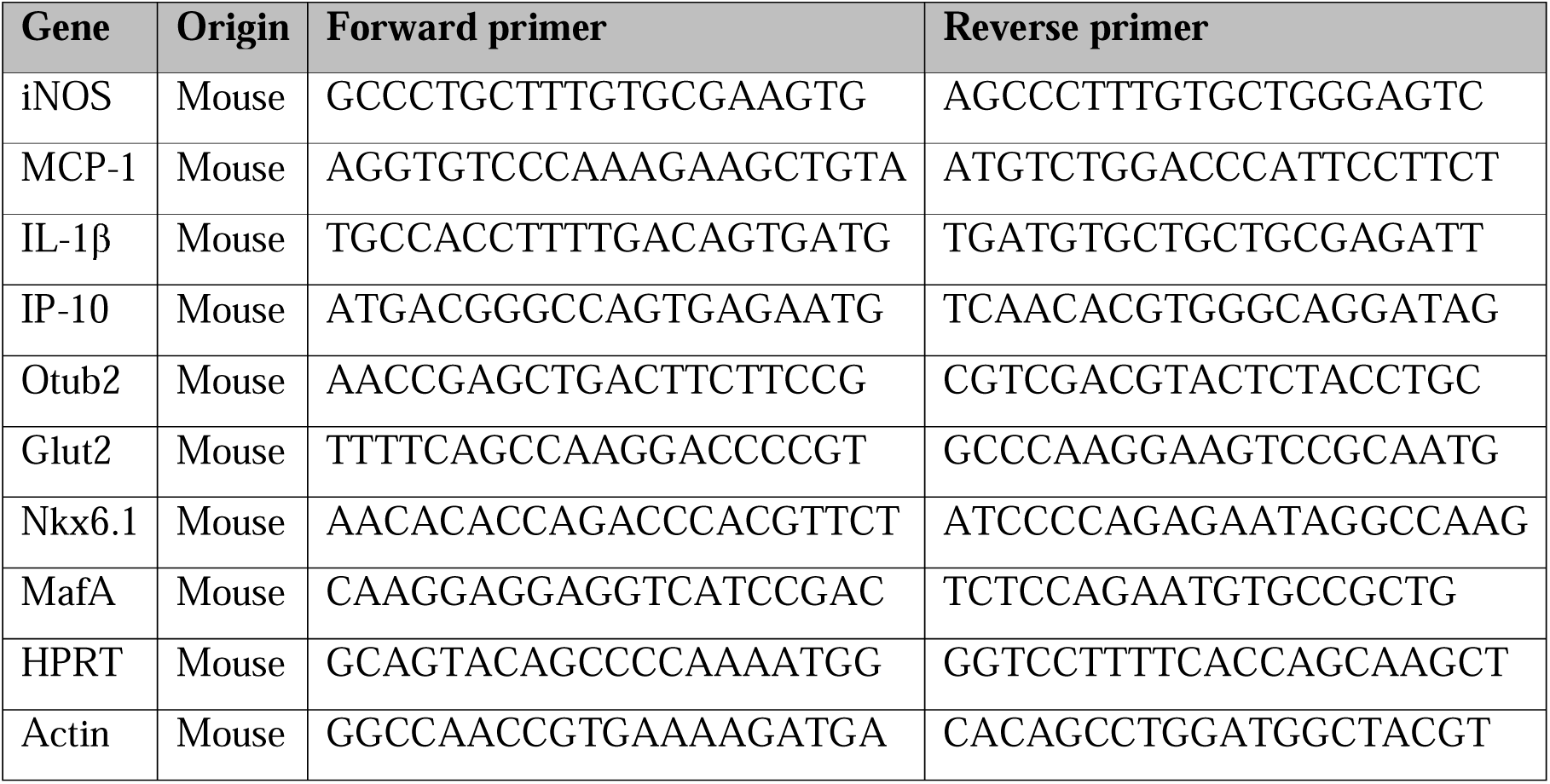
qRT-PCR primers (5’ to 3’)

### RNA sequencing

RNA was extracted from pancreas and liver of mice using the PerfectPure RNA kit (5-prime, MD) following the manufacturer’s instructions. RNA quality was assessed using Agilent 4200 TapeStation System (Agilent Technologies, Santa Clara, CA). RNA-seq libraries were generated by applying a bulk adaptation of the MARS-seq protocol, as previously described [15]. Libraries were sequenced by the Illumina Novaseq 6000 using SP mode 100 cycles kit (Illumina). Mapping of sequences to the genome and generation of the count matrix was performed by the UTAP pipeline (Weizmann Institute). Libraries normalization, filtration of low count genes and discovery of differentially expressed genes was performed using the edgeR and Limma packages in R. Gene Onthology (GO) and MsigDB pathways enrichment analysis was performed using the Camera method from the Limma package in R. The enrichment results are given in p-values, with -log10(p-value) > 1.3 considered to be significant. Functional protein association network was built using STRING (https://string-db.org/)

### Immunoprecipitation and mass spectrometry

Protein A-agarose beads were washed with ice-cold 0.1 M Tris-HCl (pH 8.0) and were incubated with Flag or GFP antibodies in 0.1 M Tris-HCl (pH 8.0) for 4 h at 4°C. Supernatants (centrifuged at 20,000xg 15 min 4°C) of cell extracts in extraction buffer, containing 0.8–1.2 mg of protein, were incubated overnight at 4°C with the immobilized antibodies. Immunocomplexes were washed three times with extraction buffer and then incubated for 30min in 4°C with PBS containing Flag or GFP peptide in order to elute Otub2 and the proteins that formed a complex with it. Immunocomplexes were mixed with Laemmle sample buffer and were resolved by 10% SDS-PAGE. The complexes were subjected to mass spectrometry analysis as described (ref – michal thesis).

### Western blot analysis

Cells were seeded and treated as indicated. Treated cells were washed three times with PBS and were harvested in buffer A (25 mM Tris-HCl [pH 7.4], 10 mM sodium orthovanadate, 10 mM pyrophosphate, 100 mM sodium fluoride, 10mM EDTA, 10 mM EGTA, and 1 mM phenylmethylsulphonyl fluoride). Cell extracts were centrifuged at 20,000xg for 15 min at 4°C, and the supernatants were collected. Samples (40–150µg) were mixed with 5x Laemmle sample buffer, boiled, and resolved by 10-12% SDS-PAGE under reducing conditions. The proteins were transferred to nitrocellulose membrane for Western blotting with the relevant antibodies.

### Generation of Otub2 knockout mice

The mouse strain C57BL/6NTac-Otub2tm1a(EUCOMM)Wtsi/WtsiH was ordered from the Wellcome Trust Sanger Institute, London, UK as part of the EUCOMM Mutant Mouse Project. One heterozygous male and two heterozygous female mice on a C57BL/6NTac background were ordered from the EMMA mouse repository. The main components of a EUCOMM vector are 5’ and 3’ homology arms that mediate homologous recombination, and a central targeting cassette that disrupts gene function and reports gene expression with a lacZ reporter. The targeting cassette is flanked by FRT recombination sites to allow removal by Flp recombinase. In addition, the vector introduces a pair of loxP recombination sites around a “critical” exon (Exons 3/4 of Otub2), which upon removal causes a frame shift, leading to complete gene inactivation. Mice were housed under standard light/dark conditions and were given access to food and water ad libitum. Experiments were approved by the Animal Care and Use Committee of the Weizmann institute of Science. For genotyping, mice tip tail genomic DNA preparations were extracted using REDExract-NAmp tissue PCR kit (Sigma), followed by amplification reactions performed with oligonucleotide pairs (see Table 2) specific for the foreign or wild-type sequences, to amplify ∼ 200-900 bp fragments.

**Table 2.**
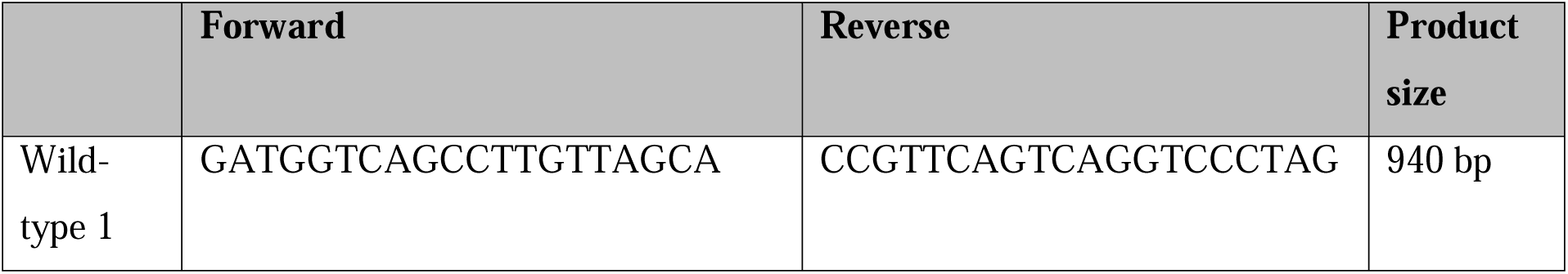
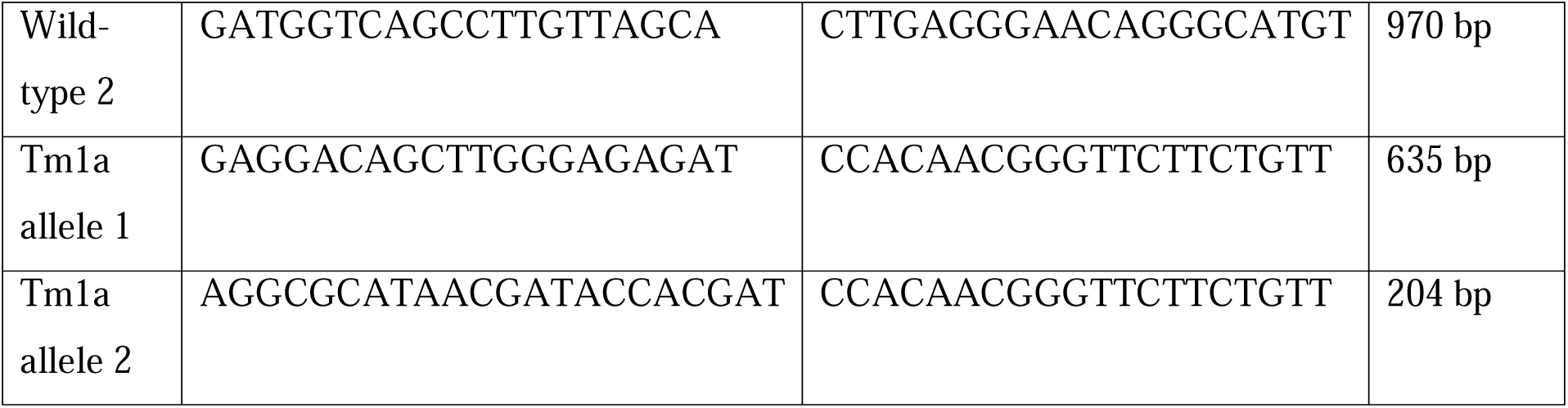
genotyping primers (5’ to 3’)

### Pancreas removal from mice

Otub2 knock-out and WT mice were sacrificed by cervical dislocation. An incision was made from the upper abdomen downwards to expose the liver and intestines. Duodenum was exposed by shifting intestines to the right, and the pancreas was isolated by pulling the intestines carefully from the duodenum downwards. Spleen was then removed and pancreas was detached from the large intestine to complete its isolation.

### Glucose tolerance test

Otub2 knock-out and WT mice (males and females, 8 weeks old) were fasted overnight with water access ad libitum. Mice were then injected intraperitoneally with glucose (1 mg/g body weight). Glucose levels were monitored using MediSense Optium Blood Glucose test strips (Abbott Laboratories, IL) on blood drawn at timed intervals from a tail vein as described [9]. Area under the curve (AUC) was done using the computeAUC function from the PharmacoGx package in R.

### Immunofluorescence of pancreatic sections

For immunofluorescence, pancreata were fixed for 24 h in 4% paraformaldehyde and then transferred to 70% ethanol until embedding in paraffin by a standard protocol (by using automated tissue processing) as follows: pancreata were first dehydrated by immersing them sequentially for 45 min. each in a series of ethanol-water mixtures (70%; 95% (x3) and 100% (x2; 30 min.)) followed by immersion with ethanol-xylene mixture (1:1; for 45 min.) and xylene (x2; 1 h). The tissues were then embedded in paraffin at 60°C (3 times for 1 h). Four-micrometer cross sections of paraffin blocks were rehydrated with xylene (x2) followed by decreasing concentrations of ethanol, microwaved in 0.01 M sodium citrate (pH 6.0; 12.5 min.) for antigen retrieval and blocking (with 0.5% Triton X 100). Sections were then incubated overnight at 4°C with antibodies targeted to specific proteins of interest as indicated. Anti-Otub2 was detected with fluorescent-tagged secondary antibodies (conjugated to Alexa-488 dye). Sections were later on washed with PBS and DAPI for nucleus staining. To minimize variability between different sections, the staining procedures for the sections were performed in parallel with the same batches of solutions. In addition, the same incubation times for fixation, permeabilization and blocking were strictly used for all processed sections.

### Statistics

Statistical analysis was performed in R. Differences between experimental conditions were determined by two-tailed Student’s t test, unless otherwise mentioned.

## RESULTS

### Over expression of Otub2 decreases NFκB activity

To characterize the effects of Otub2 on β-cells, a MIN6 cell line that stably overexpressed Flag-tagged Otub2 was generated. Over-expression of Otub2 was validated at the mRNA level by qRT-PCR (Fig. 1A) and at the protein level by Western blotting (Fig. S1A, B). Fluorescence microscopy indicated that there are no morphological changes between control Flag expressing cells and Flag-Otub2 expressing cells (Fig. S1C).

**Fig 1.**
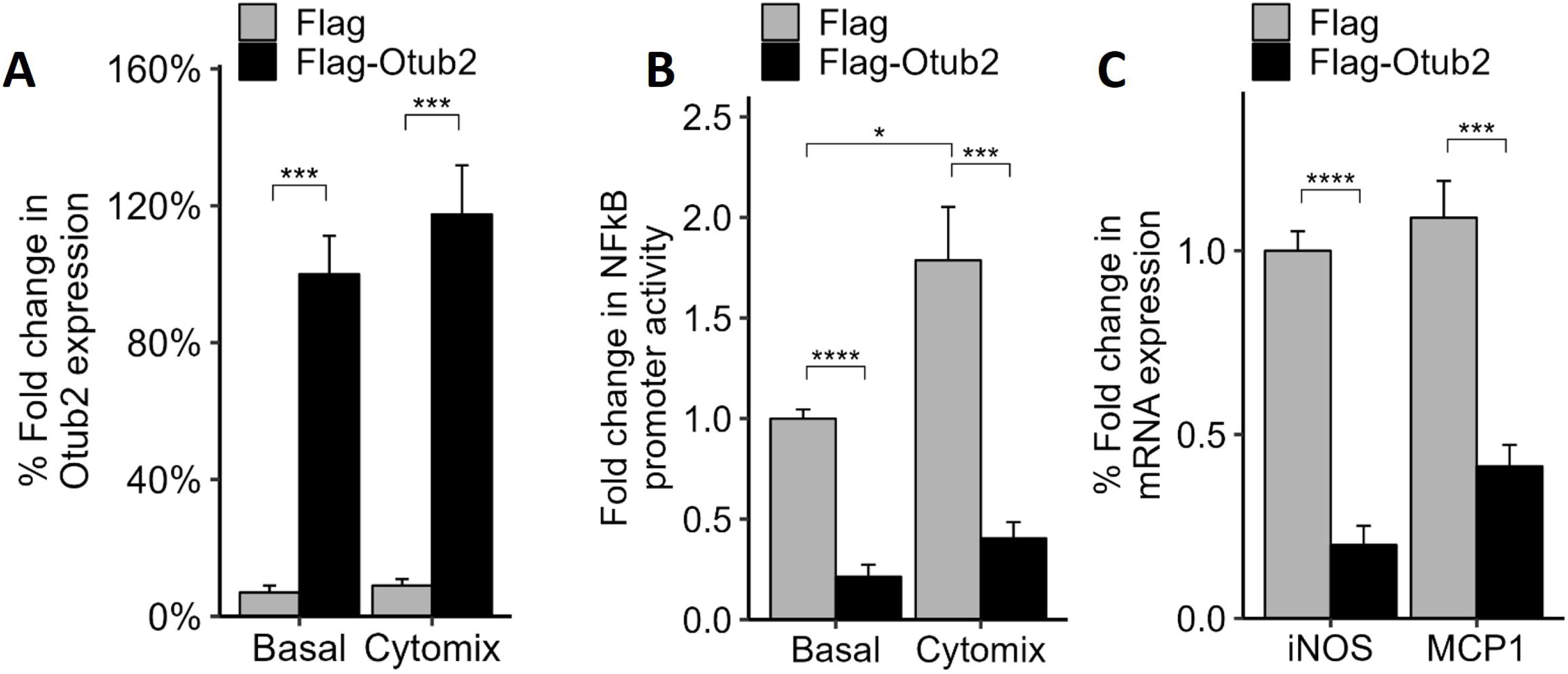
Effects of OTUB2 overexpression on NF-kB activity in MIN6 cells. **(A)** MIN6 cells were transfected with pEGFP-c1 plasmid constructs: pFlag and pFlag-Otub2 vectors. The stable cell line was subjected to qRT-PCR analysis following treatment with 1x-cytomix for 4h. Non-treated cells served as control. mRNA levels were normalized to Actin. **(B)** MIN6 cells overexpressing Flag (control) or Flag-OTUB2 were transfected with pNF-kB-MetLuc2 or with pMetLuc2 (control). 24h following transfection cells were treated with or without 1x-cytomix for 4 h. NF-kB activity was determined. Data presented were normalized to control vector transfection values. **(C)** MIN6 Flag or Flag-OTUB2 over-expressing cells were treated with 1x-cytomix for 4 h and then harvested. Total mRNA was extracted and qRT-PCR was conducted. mRNA levels of NF-kB target genes were normalized to Actin. Data are means±SEM of four replicates in each of four experiment (**B**) and of duplicates in each of three experiments (**C**). ***p<0.001 vs control Flag stable cells.

To examine the effect of Otub2 on NFκB activity, a luciferase-reporting system was employed. Min6 cells were treated with a mix of cytokines (TNFα, IL-1β IFNγ, cytomix), that significantly increase the expression of NFkB target genes [16] and promote β-cell apoptosis [16, 17]. Indeed, cytomix treatment significantly increased the level of NF-κB promoter activity by ∼1.8-fold while stable overexpression of Otub2 resulted in ∼78% decrease in NFκB promoter activity, both in control and cytomix-treated cells (Fig. 1B). Overexpression of Otub2 in cytokine-treated cells also reduced by 60% and 80%, respectively, the mRNA levels of the NFκB target genes MCP1 and iNOS (Fig. 1C). These results complement our previous findings that Otub2 knock-down increased NFκB activation that significantly increased the expression of NFκB target genes [16]. At the molecular level we could show that cytokines induced a time-dependent activation (phosphorylation) of IKK-β, the upstream activator of NFκB in control MIN6 cells. This effect was maximal following 15 min treatment with cytokines but was markedly reduced in MIN6 cells overexpressing Otub2 (Fig. S1D).

### Over-expression of OTUB2 decreases caspase 3/7 activity

The reduced NFκB activity in cells overexpressing Otub2 may induce anti-apoptotic effects. To test this possibility, we examined Caspase 3/7 activity in Otub2 over-expressing or Otub2 depleted MIN6 cells. As expected, basal Caspase 3/7 activity was significantly decreased (by ∼40%) in Otub2-overexpressing cells compared to the control Flag-overexpressing cells (Fig. 2A). In addition, the ∼5-fold increase in caspase activity observed upon cytokine treatment in control (Flag over-expressing) cells, was also significantly reduced (∼65%) in Flag-Otub2 overexpressing cells (Fig. 2A). Conversely, and consistent with our previous observations [16], silencing of Otub2 in MIN6 cells using siRNAs, significantly increased caspase 3/7 activity, both under basal conditions as well as following treatment with cytokines (Fig. 2B).

**Fig 2.**
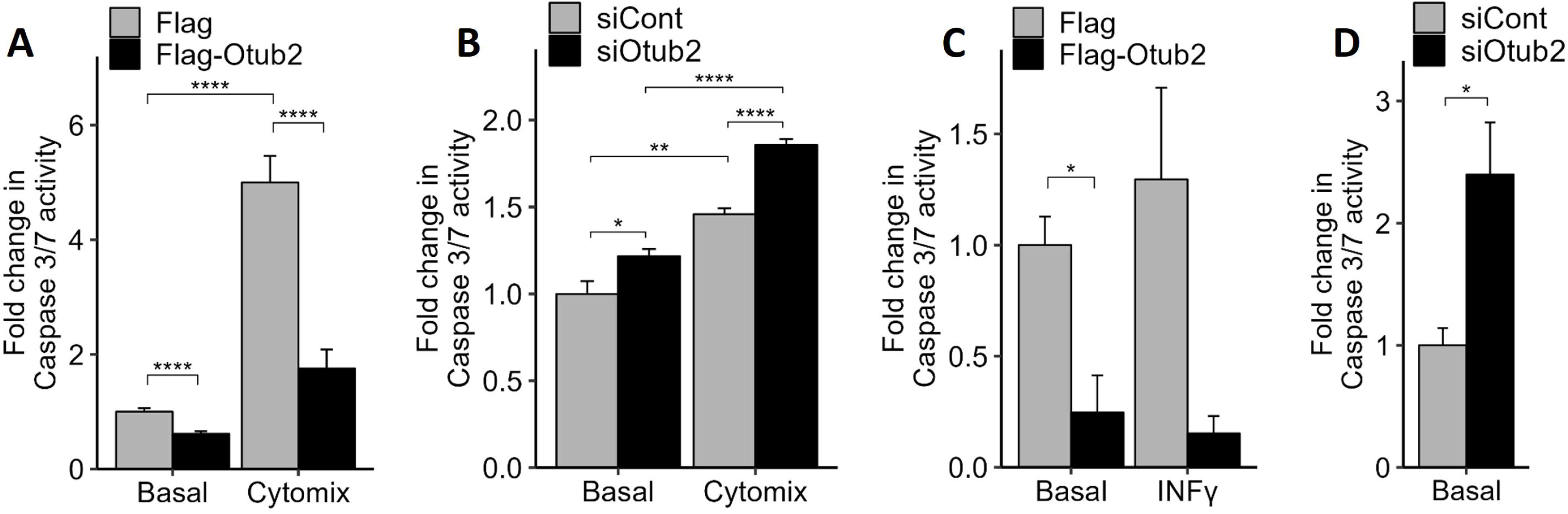
Effects of Otub2 on caspase 3/7 activity. **(A)** Control, Flag over-expressing MIN6 cells or Flag-Otub2 over-expressing cells remained untreated or were treated with 1x-cytomix for 24 h. Apoptosis was assays by caspase-3/7 activity measurements. **(B)** MIN6 cells, transfected with the indicated siRNAs remained untreated or were treated with 1x-cytomix for 24 h. Apoptosis was assays by caspase-3/7 activity measurements. Control-siRNA transfected cells served as control. **(C,D)** Dispersed human islets transiently transfected for 48 h with pFlag (control) or pFlag-Otub2 constructs, remained untreated or were treated with INFγ (C) or transfected with the indicated siRNAs (D), were assayed for apoptosis by caspase-3/7 activity. Data represent means±SEM of five replicates in each of three experiments (**A**), 4-5 replicas **(B)**, or four replicas of human islets (**C, D**) *p<0.05, ***p<0.001 vs. control cells.

Even stronger effects were observed in dispersed human islets. An approximate 5-fold decrease in caspase 3/7 activity was observed in human islets that were transfected with Flag-Otub2 construct, when compared with Flag transfected islets (Fig. 2C). Similar results were observed upon treatment of the dispersed human islets with INFγ (Fig. 2C). Conversely, opposite effects were observed in dispersed human islet in which Otub2 has been silenced by siRNA. These islets demonstrated ∼2.4-fold increase in caspase 3/7 activity compared to control, mock-transfected cells (Fig. 2D). These results indicate that over-expression of Otub2 decreases caspase 3/7 activity and, therefore, confers a pro-survival effect on human pancreatic β-cells.

### Over-expression of Otub2 affects β-cell function

To determine whether over-expression of Otub2 affects β-cell function, glucose-stimulated insulin secretion (GSIS) was studied in MIN6 cells. Basal insulin secretion was ∼2.5 higher in Flag-Otub2 expressing cells, when compared to controls (Fig. 3A). 20 mM glucose challenge increased insulin secretion 2.5-fold in control cells and 2.2-fold in Otub2 expressing cells, resulting in an approximately 2.2-fold higher GSIS in Otub2 expressing cells. These results favor our hypothesis that Otub2 promotes β-cell function, in addition to, or independent of, its inhibitory effects of NFκB activity. Next, we assessed the influence of Otub2 on Glut2, the glucose transporter mediating glucose uptake in β-cells [18]. As shown in Fig. 3B, overexpression of Otub2 dramatically increased the mRNA levels of Glut2 by approximately 20-fold. Similar increase was found in cells treated with cytokines, which by themselves drastically inhibited Glut2 expression, as previously described [19].

**Fig 3.**
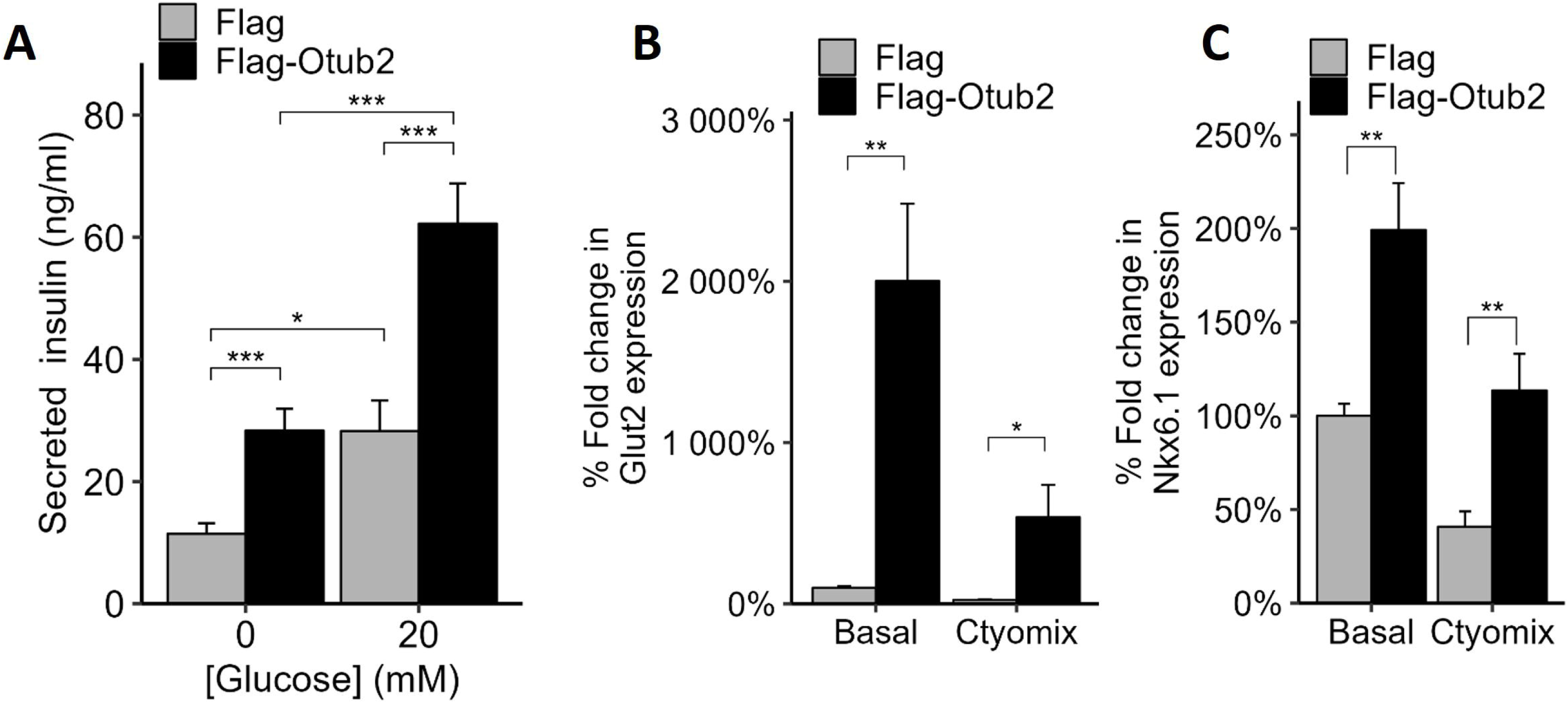
Effects of OTUB2 stable over-expression on β-cell function. (A) MIN6 cells over-expressing Flag or Flag-Otub2 were incubated for 1h in KRBH at 370C, followed by an additional 1h incubation in 0mM or 20mM glucose. Insulin secretion was then determined using insulin detecting HTRF kit (Cisbio) according to manufacturer’s instructions. (B,C) MIN6 cells over expressing Flag or Flag-Otub2 were treated with 1x-cytomix for 4h and harvested. Non-treated cells served as control. Total mRNA was extracted. Quantitative detection of GLUT2 (B) and NKX6.1 (C) was carried out by qRT-PCR. mRNA levels were normalized to Actin. Data are means±SEM of four replicates in each of four experiments (A), or two replicates in each of three experiments (B,C). *p<0.05,**p<0.01 and ***p<0.001.

Another gene of interest is Nkx6.1, a known transcription factor required for β-cell development, as well as glucose sensing and insulin secretion [20]. As shown in Fig. 3C, overexpression of Otub2 significantly increased the mRNA levels of Nkx6.1 by approximately 2- and 2.8-fold under basal or cytomix treatments, respectively. Collectively, these findings suggest that over-expression of Otub2 is involved in preserving β-cells functionality, mainly in terms of insulin secretion.

### Otub2 binding partners regulate β-cell function

To identify proteins that interact with Otub-2, immunoprecipitations (IPs) with Otub-2 antibodies were carried out using MIN6 cells transiently over-expressing Otub-2 (and appropriate controls), followed by mass spectrometry analysis. Several proteins were enriched at least 100-fold in immunoprecipitates derived from Flag-Otub2 overexpressed cells (Table 3, supplementary table S2) compared to controls. A similar trend was observed when GFP-Otub-2 Min6 cells were used. Top hits were Peg3 and Camk2d, which enhance NFκB pathway and β-cell death [21, 22]. We assume that Otub2 deubiquitinates proteins in complexes containing Peg3 and Camk2d, and thereby might inhibit propagation of the NFκB cascade that promotes β-cell apoptosis. The third protein - kv9.3, is a potassium channel responsible for repolarization of β-cell membranes after insulin release [23]. Otub2 is assumed to deubiquitinate and inhibit this channel and thereby prolong insulin release and improve β-cell function.

**Table 3.** Otub2 binding proteins (selected hits from Mass-Spec)

| Protein | Gene | Ratio<br>OTUB2/ FLAG | Ratio<br>OTUB2/GFP | Function |
| --- | --- | --- | --- | --- |
| OTUB2 | Otub2 | 145.09 | 179.16 |  |
| PEG3 | Peg3 | Inf | Inf | Acts synergistically with TRAF2 through activation of NFκB |
| KCC2D | Camk2d | Inf | 133.39 | Calcium/calmodulin-dependent kinase |
| KCNS3 | Kcns3 | Inf | 29.11 | Voltage-gated potassium channel subunit Kv9.3 |
| SSRD | Ssr4 | Inf | 8.37 | Binds clacium to the ER membrane and regulates the retention of ER resident proteins |
| SSRG | Ssr3 | 2.53 | 3.35 | Binds clacium to the ER membrane and regulates the retention of ER resident proteins |
| AT1A1 | Atp1a1 | Inf | 3.62 | Sodium/potassium-transporting ATPase |
| ATPG | Atp5c1 | Inf | 2.25 | Mitochondrial membrane ATP synthase |
| AT2A3 | Atp2a3 | 3.09 | 4.56 | Catalyzes the hydrolysis of ATP coupled with the transport of calcium |

### Otub-2 affects β-cell functionality in-vivo

The effects of Otub-2 on β-cell functionality *in-vivo* were evaluated next. To this end, we employed a mouse model that harbors either heterozygous (Otub2^+/-^) or homozygous (Otub2^-/-^) whole-body deletion of the Otub2 gene. Pancreata were isolated from wild-type (WT, Otub2^+/+^) and heterozygous Otub2 knockout mice (het, Otub2^+/-^), and the expression of several NFκB target genes was evaluated. As shown in Fig. 4A, the expression levels of Otub2 itself were decreased by approximately 2-fold in Otub2^+/-^ mice compared to Otub2^+/+^ mice, confirming the expected partial down-regulation of the Otub2 gene expression in the Otub2^+/-^ animals. Conversely, there was a marked increase in the expression of several NFκB targets in Otub2^+/-^ mice, including IP-10, MCP-1 and IL1-β, whose expression levels were increased by approximately 5-, 6- and 2-fold, respectively (Fig. 4A). These results suggest an increased inflammatory state of the pancreas upon partial Otub2 knockout, supporting the physiological role of Otub2 as a negative regulator of NFκB activity in pancreatic islets *in vivo*. Immunofluorescence staining of pancreatic sections from those mice, confirmed the expected decrease in Otub2 protein levels in the Otub2^+/-^ and ^-/-^ mice (Fig. 4B). Next, glucose tolerance tests (GTT) were performed in Otub2^+/+^, Otub2^+/-^ and Otub2^-/-^ mice.

**Fig 4.**
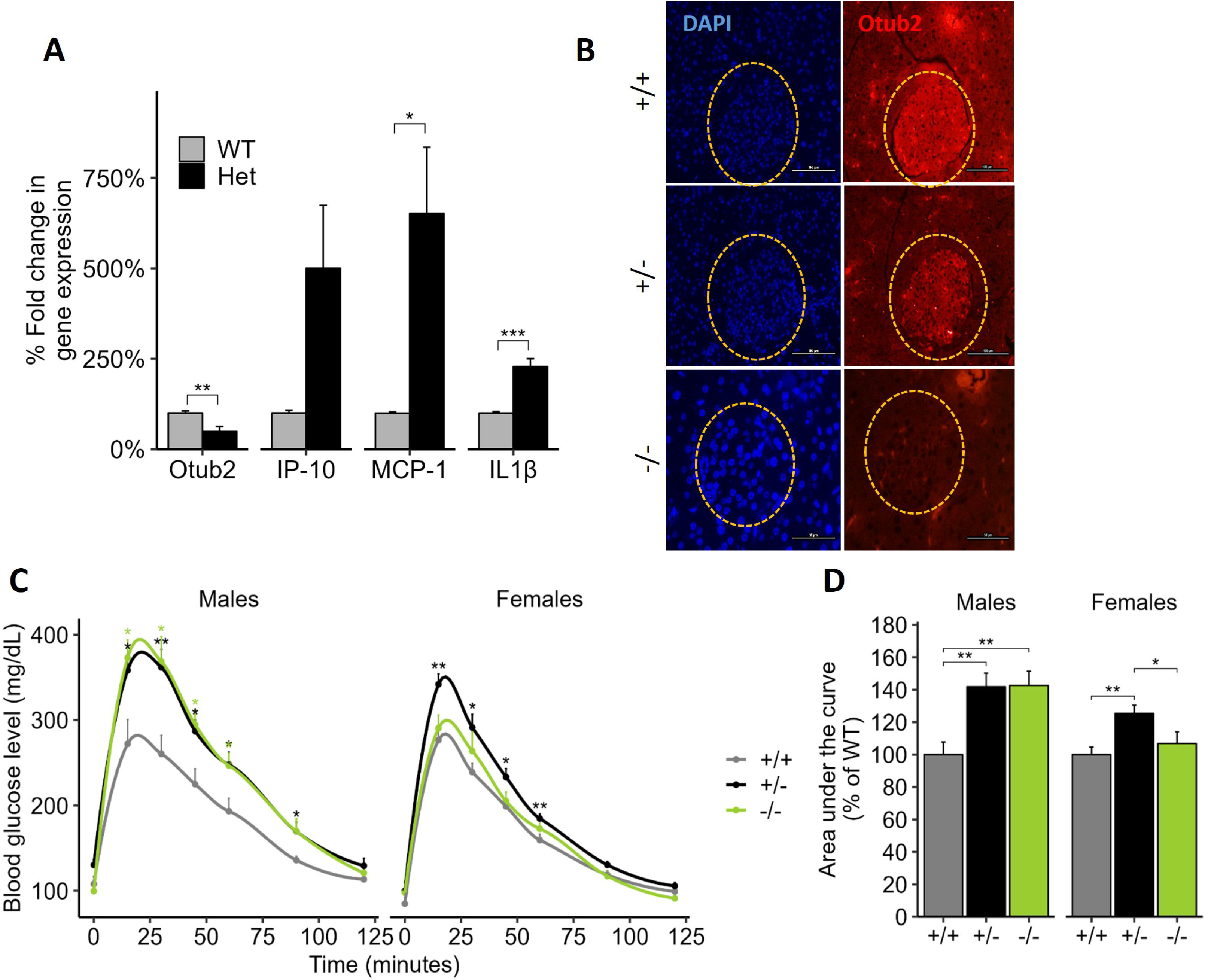
Effects of OTUB2 knock-down on pancreatic NF-kB activity in vivo. Pancreata from Otub2^+/+^ (WT) and Otub2^+/-^ (het) female and male mice were excised. **(A)** Total mRNA was extracted and qRT-PCR was conducted. mRNA levels of NF-kB target genes were normalized to Actin. Data is Otub2^+/+^ n=5; Otub2^+/-^ n=5. Data represent means±SEM of three experiments *p<0.05,**p<0.01 and ***p<0.001 vs. Otub2^+/+^ mice. **(B)** Pancreas sections were stained for OTUB2 and DAPI. **(C-D)** *Otub2 knock-out mice are glucose intolerant*. Male and female mice at 8 weeks of age were subjected to intra-peritoneal (i.p.) glucose tolerance test (GTT; 1.8 g D-glucose per kg body) after over-night fasting. Blood samples were taken at the indicated time points (0’-120’) and glucose levels were determined by a glucometer (**C).** A graph representing the area under the curve of the GTT is indicated as (**D**). Data represent means±SEM of six experiments *p<0.05,**p<0.01 vs. Otub2^+/+^ mice. Males: Otub2^+/+^-n=7; Otub2^+/-^-n=12; Otub2^-/-^-n=8. Females: Otub2^+/+^ n=8; Otub2^+/-^ n=10; Otub2^-/-^ n=8.

As shown in Fig. 4C, basal glucose levels of Otub2^+/+^, Otub2^+/-^ and Otub2^-/-^ male and female mice were approximately similar. However, blood glucose levels, at all-time points tested, were significantly higher in Otub2^+/-^ and Otub2^-/-^ male mice, when compared to Otub2^+/+^ animals (Fig. 4C). Accordingly, the area under the curve (AUC) was ∼40% greater in Otub2^+/-^ and Otub2^-/-^ male mice compared to WT controls (Fig. 4D). The effects of Otub2 deletion on the female mice were a bit more complex. There were no differences in GTT between Otub2^+/+^ and Otub2^-/-^ female mice, but the Otub2^+/-^ animals exhibited a slight, yet significant higher GTT response (Fig. 4C) that was also evident by a modest, yet significant higher (approximately 20%) AUC (Fig. 4D). These results suggest a positive role for Otub2 in the improvement of islet functionality under physiological conditions in an *in-vivo* setting, with possible compensatory pathways in KO female mice.

### Deletion of Otub2 affects pancreatic gene expression patterns

To reveal the genes whose expression is affected upon Otub-2 deletion, RNA was extracted from pancreases of Otub2^-/-^, Otub2^-/+^, and Otub2^+/+^ (WT) mice and RNAseq analysis was performed. The expression levels of Otub2 were indeed below detection level in most Otub2^-/-^ and Otub2^-/-^ pancreases (Fig. S2A). Both heterozygous and homozygous pancreases showed significant different gene expression patterns compared to WT pancreases, as shown in the volcano plots (Fig. S2B). Of note, at least 24 and 23 genes were significantly up- or down regulated, respectively, in both heterozygous and homozygous vs. WT mice (Supplemental Table S1). Enrichment analysis of curated signatures was performed, employing the MsigDB database and using the Camera method [24] (Fig. S2C). Several gene sets were significantly upregulated. Most significant were the MOOTHA_VOXPHOS genes [25], involved in oxidative phosphorylation. Other relevant families that supported our experimental findings were WANG_NFKB_TARGETS pathway [26] that includes NFκB target genes; REACTOME_DIABETES_TARGETS [27], and ALCALA_APOPTOSIS [28]; all gene sets expected to be upregulated upon depletion of Otub2.

To gain a deeper insight into the gene networks affected by Otub-2 depletion, pathway enrichment analysis was carried out using the gene onthology (GO) database. Such analysis revealed that potassium ion transport was significantly affected in Otub-2 ^-/-^ and Otub-2 ^-/+^ mice (Fig. 5A, B), with several genes related to this pathway being down-regulated in OTUB2 KO animals (Fig. 5B). One is Ank2 that encodes ankyrin-2 (AnkB), which is essential for localization and membrane stabilization of potassium channels. Another relevant gene is KCNC3, encoding a voltage-dependent potassium channel involved in insulin secretion [29], and Cacna1a encoding the pore-forming α1A subunit of the neuronal channel P/Q, which belongs to the superfamily of voltage-gated calcium channels.

**Fig 5.**
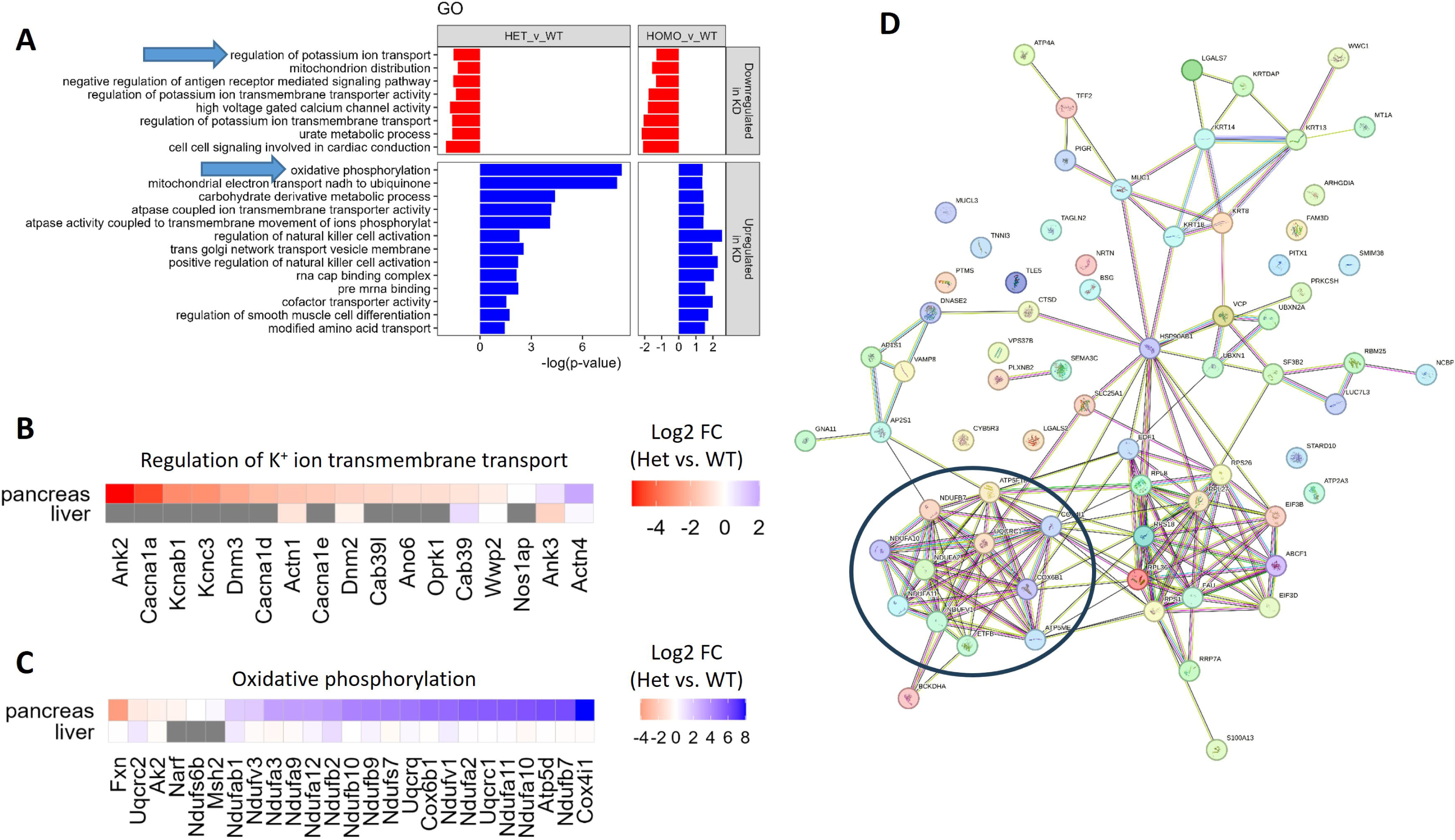
RNA-seq analysis of pancreas from OTUB2 KO mice. Pancreases and livers were extracted from OTUB2^+/+^ (WT, n = 3), OTUB2^+/-^ (Heterozygotes, n = 5) and OTUB2^-/-^ (Homozygote, n = 3) mice. RNA was extracted and RNA-seq was performed to determine differential gene expression between the homozygotes/heterozygotes and the WT mice. **(A)** Geneset enrichment of gene onthology (GO) terms, performed by the CAMERA method. Arrows indicate interesting GO terms. Analysis is based on gene expression in the pancreases. **(B,C)** Heatmaps representing the fold change (log2) in gene expression between heterozygotes and WT mice in pancreases and livers. Genes related to potassium ion transport (B) and oxidative phosphorylation (B) are taken from the respective GO terms. (D) STRING analysis of genes significantly upregulated in OTUB2^+/-^ vs. OTUB2^+/+^ pancreases. The black circle depicts cluster of genes associated with oxidative phosphorylation.

Conversely, gene-enrichment analysis revealed that the gene family encoding oxidative phosphorylation named MOOTHA_VOXPHOS [25] is mostly upregulated in heterozygous and homozygous OTUB2 KO mice (Fig. 5A, 5C). This gene family is also upregulated when the MsigDB database is analyzed according to the Camera method (Fig. S2C, S2D). Finally, STRING analysis of the genes significantly upregulated in the Otub-2 ^-/+^ mice has revealed a cluster of oxidative phosphorylation related genes in the pancreas of these mice (Fig. 5D).

Given that β-cells are highly susceptible to oxidative stress [30, 31], the increase in oxidative phosphorylation might be associated with β-cells dysfunction and apoptosis observed upon OTUB-2 depletion. Of note, the ATP dependent potassium channels, discussed above, are specific targets of oxidative damage [32]. The upregulated genes in OTUB2 KO pancreases, related to oxidative phosphorylation, are almost exclusively encoding for components of the electron transport chain (ETC) complexes in the mitochondria (Fig. 5C). The highest upregulated gene in this pathway, Cox4i1, was previously shown to be upregulated in type 2 diabetic mice [33]. This suggests that oxidative stress is a possible mechanism by which knockdown of OTUB-2 impairs β-cells function.

In general, alterations in the transcriptomic landscape of pancreata depleted of Otub-2 are quite unique, as largely different alterations were observed when comparing livers of heterozygous and homozygous OTUB2 KO mice to livers of WT animals (Fig. S3A, S2B). Still, pathway enrichment analysis revealed that oxidative phosphorylation genes, as well as components of the ETC complexes in the mitochondria, are also enriched in the liver (Fig. S3C). Although these genes are not upregulated to the same extent as they do in the pancreas (Fig. 5C, S2D), our results suggest that knockdown of OTUB-2 ubiquitously upregulates genes involved in oxidative phosphorylation and mitochondrial ETC.

## Discussion

In our previous studies we identified OTUB2 as a pro-survival ’hit’ in siRNA screens using primary human islets and MIN6 cells [9], implicating Otub2 as a potential physiological regulator of β-cell survival. We previously showed that down-regulation of Otub2 by siRNAs in MIN6 cells and human islets, increased caspase-3/7 activity, reduced GSIS and elevated expression of NFκB target genes, both under basal levels and following cytokine treatment [9]. In this study, we employed animal models to strengthen these findings and reveal new aspects of Otub2 action as an inhibitor of NFκB signaling and as a pro-survival protein of human β-cells that promotes β-cells functionality.

Several lines of evidence support this conclusion. First, we showed that over-expression of Otub2 decreases NFκB activity and expression of its target genes MCP-1 and iNOS. The increased expression of these genes was also observed in pancreatic sections derived from Otub2-knockout mice, thus highlighting the physiological relevance of these findings. Given that NFκB signaling promotes apoptosis of pancreatic β-cells [34, 35], it was of no surprise that overexpression of Otub2 inhibited cytokine production and Caspase 3/7 activity in β-cell lines and human pancreatic islets, while inhibition of Otub2 expression exerted opposite effects.

At the molecular level, earlier work has shown that the extent of ubiquitination of TRAF6, a key player in cytokine-induced NFκB activation, is elevated in MIN6 cells upon Otub2 silencing [9, 36]. Additional studies showed that Otub2 is a negative regulator of type I IFN induction through deubiquitination of TRAF3 and TRAF6 [37]. Thus, our working hypothesis predicts that K63-deubiquitination of TRAF6, induced by Otub2, inhibits cytokine-induced K48-ubiquitinqtion and degradation of IkB. This inhibits propagation of NFκB signaling and the expression of NFκB target genes such as MCP-1, IP-10 IL-1β and iNOS [38, 39], that contribute to β-cell demise (Fig. 6). Indeed, we could show that activation (phosphorylation) of IKK-β, the upstream activator of NFκB is markedly inhibited upon overexpression of Otub2 in β-cells. Of greater physiological relevance are the effects of Otub2 on β-cell function. We showed that overexpression of Otub2 promotes insulin secretion in cultured β-cell lines, whereas partial or complete knockdown of the Otub2 gene, mainly in male mice, significantly impairs their glucose tolerance. Interestingly, while depletion of Otub2 in heterozygous mice resulted in impaired GTT, in homozygous mice the effect was evident only in male mice, suggesting, a yet unknown, gender-specific compensatory effect that affects only female homozygous knockout animals.

**Fig. 6.**
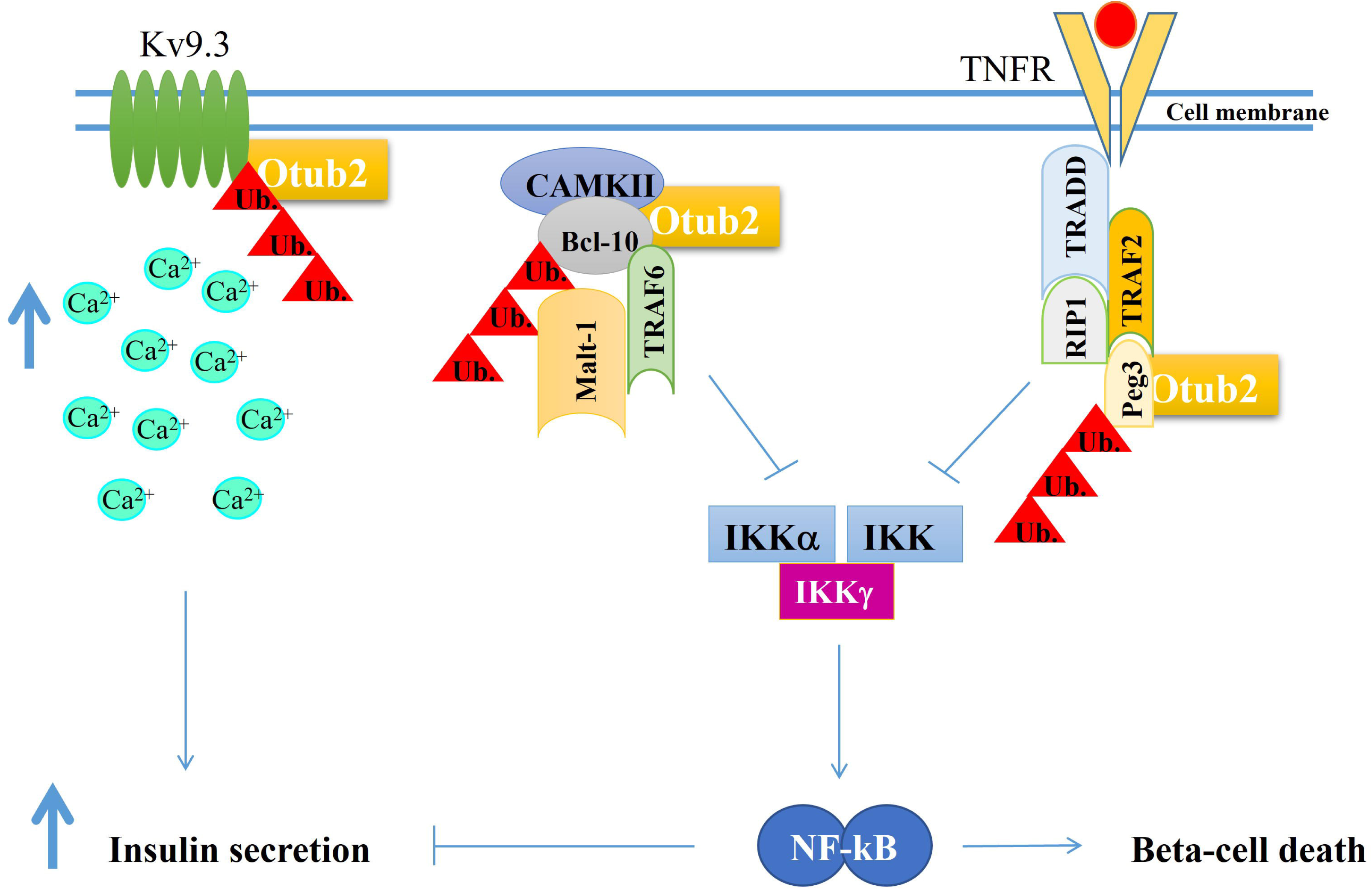
Model for Otub2 potential binding partners in β cells. Analysis of mass spectrometry results indicate of more potential binding partners involved in the NF-kB pathway (e.g Peg3 and Bcl-10) Otub2 presumably deubiquitinates these proteins which eventually inhibits β-cell apoptosis. Moreover, Otub2 potentially binds and inhibits Kv9.3 voltage-gated potassium channel, which helps prolong GSIS by delaying repolarization of the cell’s membrane.

Several mechanisms could account for the impaired glucose tolerance in Otub2^+/-^ and Otub2^-/-^ animals. Most likely, these effects could be attributed to the inhibitory effects of Otub2 on NFκB activity as discussed above. We found that Otub2 forms complexes with Peg3 and Camk2d, both enhancer of NFκB pathway and β-cells death [21, 22]. Peg3, a regulator of the TNF response, acts synergistically with TRAF2 to activate the NFκB signaling pathway [21]. Hence, deubiquitination of Peg3 by Otub2, might inhibit NFκB signaling in response to cytokines. Another key player in this process might be the calcium/calmodulin-dependent kinase (CaMKIID) that forms complexes with Otub2. CaMKIID phosphorylates Bcl-10 during activation of the NFκB pathway. Ubiquitination of phospho-Bc1-10 promotes its interaction with Malt-1, which binds TRAF6 [40], another Otub2 target [9]. Hence, complex formation between Otub2 and CaMKIID, might facilitate Bcl-10-CaMKIID interaction and prevents its ubiquitination by TRAF6 and termination of NFκB signaling.

Yet, Otub2 seems to exert additional effects that might be only partially related to NFκB signaling. Overexpression of Otub2 significantly increases expression of Glut2, the main glucose transporter of β−cells, which is essential for glucose-stimulated insulin secretion [18]. Further, increased expression of Otub2 upregulates the expression of Nkx6.1, a transcription factor essential for maintaining the functional and molecular traits of mature β-cells (e.g. insulin biosynthesis and insulin secretion [41]).

A broader perspective concerning the physiological functions of Otub2, was gained by analyzing alterations in the transcriptomic landscape of pancreata depleted of Otub-2. First, it is obvious that the effects of Otub2 on the transcriptomic landscape seems to be specific as different gene sets are affected when we compare liver to pancreata in wild type vs. OTUB2 knockout animals.

Concerning the pancreas, the analysis highlighted a significant reduced expression of genes that regulate potassium ion channel transport, which directly influences insulin secretion [42]. Of interest, one of this family members, Kv9.3 that is expressed in human pancreatic islets [23], forms complexes with Otub2. This channel is involved in repolarization of excitable cells, therefore blocking the activity of this delayed rectifier potassium channel is expected to increase intracellular free calcium and promote GSIS [23]. Potassium channels need to be closed for proper insulin secretion, in a process mediated by ATP [42]. Therefore, activating mutations in potassium channels may induce adult and neonatal diabetes [43]. Indeed, impaired expression of a potassium channel regulator AnkB, that occurs in OTUB2 KO mice, impairs insulin secretion and induces diabetes [44]. Similarly, downregulation of the dynamin genes DNM2 and DNM3 [45], observed in OTUB2 KO mice, attenuates internalization of potassium channels on one hand, while inhibiting exocytosis of insulin granules [46, 47]. Hence, maintenance of higher concentrations of potassium channels at the plasma membrane of OTUB2 KO mice, might impair glucose-stimulated insulin secretion which is the characteristic feature of these animals.

In line with this hypothesis, knockdown of Otub2 promotes transcription of oxidative phosphorylation pathways, with the NDUF gene-family members being the targets. These proteins are component of complex I in the mitochondrial electron transport chain [48]. Hence, increased ATP production in the KO animals might be a compensatory mechanism utilized by these mice to foster closure of the highly abundant potassium channels, and thus promote insulin secretion. This, on the other hand, could also induce oxidative stress which is detrimental for β-cell function and survival [20]. It is well established that β-cells are sensitive to oxidative stress due to their low antioxidative capacity [30]. Otub2 silencing in ovarian cancer stabilizes sorting nexin 29 pseudogene 2 (SNX29P2), which prevents hypoxia-inducible factor-1 alpha (HIF-1α) from von Hippel-Lindau tumor suppressor-mediated degradation. Elevated HIF-1α activates the transcription of carbonic anhydrase 9 and drives ovarian cancer progression and chemoresistance by promoting glycolysis [49]. Hence, reprogramming and activation of ATP-generating systems might be a common outcome of Otub2 silencing. Yet, Otub2 silencing might exert opposite effects in some model systems. For example, OTUB2 depletion reduces glucose consumption, lactate production, and cellular ATP production in colorectal cancer cells [50]. Similarly, decreased expression of MOOTHA_VOXPHOS genes contributes to the development of type-2 diabetes in human muscles [25]. Hence, the effects of Otub2 on oxidative phosphorylation and ATP production might well be tissue specific. The wide differences in gene expression profiles between liver and pancreas of OTUB2 KO mice support this conclusion.

In summary, our findings highlight the role of Otub2 as an anti-apoptotic regulator of β-cell survival that preserves their mass and functionality. Otub2 inhibits NFκB signaling and cytokine production, downregulates membranal potassium channels; promotes insulin secretion and reduces oxidative stress. This raises the possibility of developing novel therapeutic strategies that target Otub2 and increase its expression for the benefit of diabetic patients.

## Acknowledgements

We thank Dr. Eythan Elhanany for insightful comments and discussions. Human islets were provided through the JDRF award 31-2008-413 (ECIT Islet for Basic Research program).

## Funding

This work was supported by grants from the Juvenile Diabetes Research Foundation International (17-2013-442) and the Israel Science Foundation (759/09).

## Duality of interest statement

The authors declare that there is no duality of interest associated with this manuscript.

## Contribution Statement

M.A., R.I., S. B-H., and Y. V. performed the studies and analyzed the data. S.S., S.L, Y.V, and Y.Z. contributed to the design of the experiments and drafting the manuscript. All authors gave final approval of the version to be published.

## Notes

### Competing Interest Statement

The authors have declared no competing interest.

