## Supplemental figures and tables for "The Deubiquitinating Enzyme Otub2 Modulates Pancreatic Beta-Cells Function and Survival"

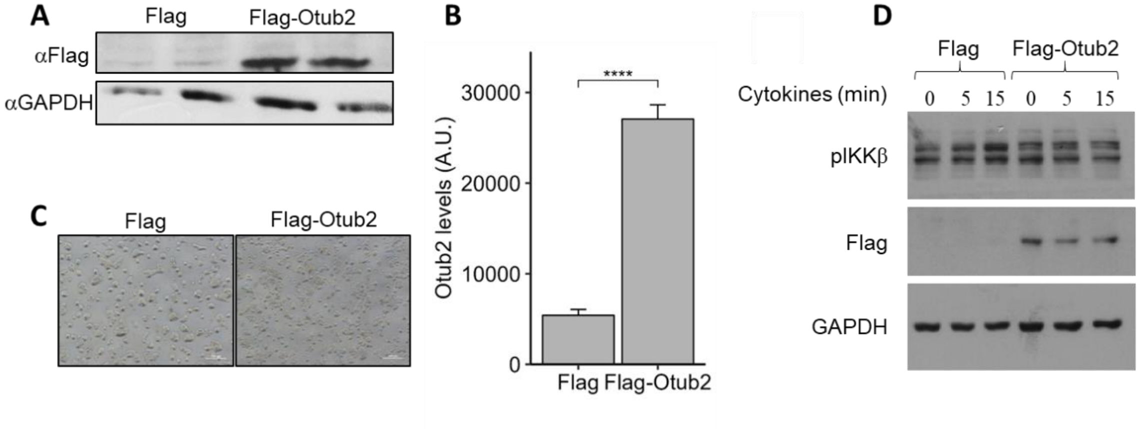


**Fig. S1 -** ***Otub2 expression levels in MIN6 stable cell line*.** **(A)** Flag and Flag-OTUB2 overexpressing cells were used for protein extraction, resolved by SDS-PAGE and western blotted using anti-Flag antibody. Protein levels were normalized to GAPDH or to Tubulin. **(B)** Densitometry analysis was used to quantify OTUB2 protein levels. **(C)** After selection process, Flag and Flag-Otub2 overexpressing cells were photographed using florescence microscope. **(D)** MIN6 cells overexpressing Flag (control) or Flag-OTUB2 were treated with 1x-cytomix for 5 or 15 minutes, or were left untreated. Cells were then harvested, proteins were extracted and resolved by SDS-PAGE. Western blot analysis was conducted with anti-phospho-IKK and anti-Flag antibodies. Protein levels were normalized to GAPDH levels. Data represent means±SEM of two experiments in duplicates and triplicates (A,B) or of three independent experiments in duplicates (C); ***p<0.001 vs control cells.

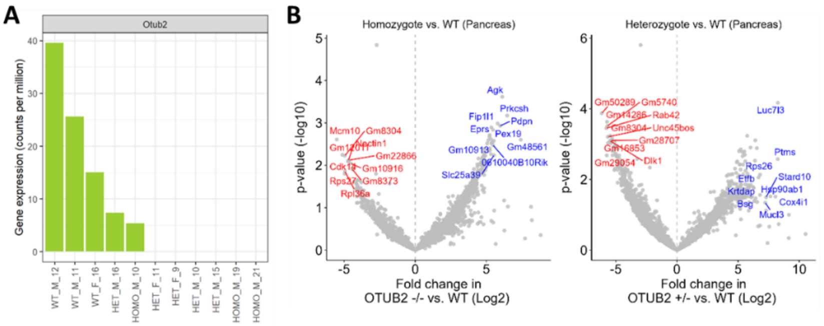


C

| Pathway | OTUB2 +/- vs. WT |
|---|---|
| MOOTHA_VOXPHOS | 8.83 |
| FRASOR_TAMOXIFEN_RESPONSE_UP | 2.76 |
| REACTOME_DIABETES_PATHWAYS | 2.74 |
| WANG_NFKB_TARGETS | 2.38 |
| ALCALA_APOPTOSIS | 2.21 |
| GRAESSMANN_APOPTOSIS_BY_DOXORUBICIN_UP | 2.02 |

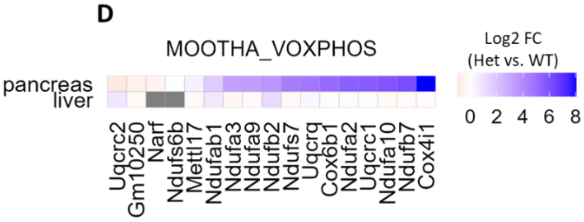


**Fig S2 – *RNA-seq analysis of pancreases from OTUB2 KO mice*.** Pancreases were extracted from OTUB2$^{+/+}$ (WT), OTUB2$^{+/-}$ (Heterozygotes) and OTUB2$^{-/-}$ (Homozygote) mice. RNA was extracted and RNA-seq was performed. (A) Expression of OTUB2 in the pancreases, as determined by the RNAseq. (B) Volcano plot showing the fold changes in genes expression between homozygotes (left) or heterozygotes (right) vs. WT pancreases. (C) Geneset enrichment analysis using the MsigDB signature set, performed using the CAMERA method. The numbers indicate the significance of the enrichment (in terms of –log10[p-value]) between heterozygotes and WT mice. (D) Heatmap representing the fold change (log2) in gene expression between heterozygotes and WT mice in pancreases and livers. Genes are taken from the MOOTHA_VOXPHOS geneset.

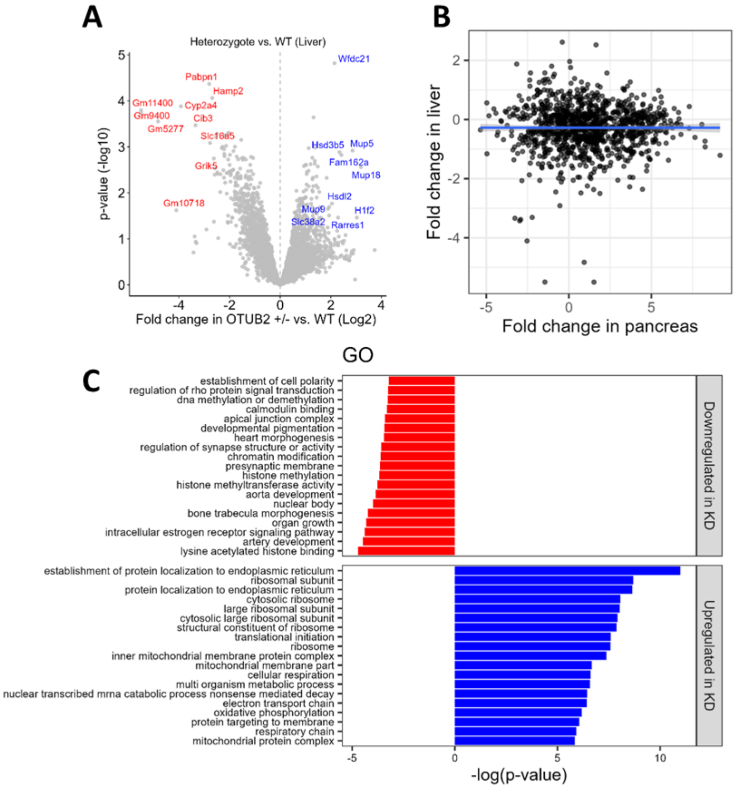


**Fig S3 – *RNA-seq analysis of livers from OTUB2 KO mice*. L**ivers were extracted from $OTUB2^{+/+}$ (WT) and $OTUB2^{+/-}$ (Heterozygotes) mice. RNA was extracted and RNA-seq was performed. **(A)** Volcano plot depicting fold change in gene expression between heterozygotes and WT livers. **(B)** Comparisons between the fold change in each gene expression between heterozygotes and WT pancreases (in the x-axis) and livers (in the y-axis). **(C)** Geneset enrichment of gene onthology (GO) terms, performed by the CAMERA method, using the RNA-seq results of the livers.

### Supplemental tables

Table S1 – Genes significantly up or down regulated in both homozygotes and heterozygotes vs. WT

| | Homozygotes vs WT | | Heterozygotes vs WT | |
|---|---|---|---|---|
| genes | Fold changes (log2) | p-values | Fold changes (log2) | p-values |
| PRKCSH | 6.46 | 0.00 | 5.60 | 0.00 |
| PDPN | 5.97 | 0.00 | 3.73 | 0.04 |
| FIP1L1 | 5.79 | 0.00 | 4.54 | 0.01 |
| CLTC | 5.31 | 0.00 | 4.19 | 0.01 |
| NCBP3 | 4.92 | 0.01 | 5.08 | 0.00 |
| NDUFB9 | 4.84 | 0.03 | 4.48 | 0.04 |
| LUC7L3 | 4.83 | 0.04 | 8.26 | 0.00 |
| MERTK | 4.82 | 0.01 | 3.72 | 0.04 |
| RRP7A | 4.79 | 0.01 | 5.37 | 0.00 |
| WDR37 | 4.73 | 0.04 | 4.52 | 0.03 |
| EPB41 | 4.67 | 0.03 | 4.33 | 0.03 |
| GM6129 | 4.43 | 0.05 | 4.15 | 0.05 |
| SLBP | 4.33 | 0.03 | 3.60 | 0.05 |
| GM15620 | 4.21 | 0.03 | 3.97 | 0.03 |
| LARP4B | 4.20 | 0.03 | 3.81 | 0.04 |
| GM26244 | 4.14 | 0.02 | 5.22 | 0.00 |
| SLC34A2 | 4.12 | 0.01 | 3.28 | 0.04 |
| 1700017B05RIK | 4.11 | 0.02 | 3.37 | 0.04 |
| CTBP2 | 4.08 | 0.04 | 3.57 | 0.05 |
| NRGN | 4.07 | 0.03 | 4.28 | 0.02 |
| CHERP | 4.07 | 0.04 | 4.12 | 0.02 |
| SNX21 | 4.00 | 0.03 | 3.57 | 0.04 |
| FAM228A | 3.97 | 0.03 | 4.40 | 0.01 |
| 4932416K20RIK | 3.51 | 0.04 | 3.42 | 0.03 |
| GM10801 | -1.80 | 0.03 | -2.22 | 0.00 |
| GM10717 | -2.56 | 0.05 | -2.92 | 0.01 |
| GM10800 | -2.71 | 0.00 | -2.98 | 0.00 |
| GM21738 | -2.76 | 0.02 | -3.26 | 0.00 |
| PAQR3 | -3.38 | 0.05 | -4.42 | 0.00 |
| GM4319 | -3.74 | 0.05 | -4.19 | 0.01 |
| A530041M06RIK | -3.79 | 0.03 | -4.02 | 0.01 |
| GM47592 | -3.83 | 0.05 | -4.39 | 0.01 |
| GM28707 | -3.84 | 0.03 | -5.42 | 0.00 |
| GM14286 | -3.88 | 0.02 | -5.72 | 0.00 |
| A430033K04RIK | -4.04 | 0.02 | -3.86 | 0.01 |
| SLC2A9 | -4.05 | 0.03 | -4.34 | 0.01 |
| HEATR5A | -4.10 | 0.02 | -4.30 | 0.01 |
| ST3GAL2 | -4.13 | 0.03 | -3.37 | 0.04 |
| GM16105 | -4.17 | 0.04 | -4.37 | 0.01 |
| FAIM2 | -4.37 | 0.01 | -2.95 | 0.05 |
| GM8373 | -4.44 | 0.01 | -4.49 | 0.00 |
| GM10916 | -4.52 | 0.01 | -4.88 | 0.00 |
| GM22866 | -4.65 | 0.01 | -5.19 | 0.00 |
| CDK14 | -4.73 | 0.01 | -3.23 | 0.04 |
| GM8304 | -4.95 | 0.01 | -5.46 | 0.00 |
| GM12011 | -5.13 | 0.01 | -3.80 | 0.02 |
| MCM10 | -5.52 | 0.00 | -4.97 | 0.00 |

Table S2 – Full results of Otub2 binding genes

| Protein | Gene Name | Ratio OTUB2/Cont (FLAG) | Ratio OTUB2/Cont (GFP) | FLAGotub2 | FLAG | GFPotub2 | GFP | # peptides | Seq. Coverage |
|---|---|---|---|---|---|---|---|---|---|
| sp\|Q11136\|PEPD_MOUSE | Pepd | Inf | Inf | 6.37E+06 | 0.00E+00 | 1.16E+08 | 0.00E+00 | 1 | 1.8 |
| sp\|Q3URU2\|PEG3_MOUSE | Peg3 | Inf | Inf | 2.40E+07 | 0.00E+00 | 1.30E+08 | 0.00E+00 | 1 | 1.5 |
| sp\|Q6PHZ2\|KCC2D_MOUSE | Camk2d | Inf | 133.39 | 2.19E+08 | 0.00E+00 | 1.73E+09 | 1.30E+07 | 3 | 10.6 |
| sp\|Q60932\|VDAC1_MOUSE | Vdac1 | Inf | 47.47 | 5.96E+06 | 0.00E+00 | 1.35E+09 | 2.85E+07 | 2 | 10.8 |
| sp\|Q8BQZ8\|KCNS3_MOUSE | Kcns3 | Inf | 29.11 | 5.76E+08 | 0.00E+00 | 1.11E+08 | 3.80E+06 | 1 | 1.4 |
| sp\|Q62318\|TIF1B_MOUSE | Trim28 | Inf | 15.92 | 8.82E+07 | 0.00E+00 | 2.88E+08 | 1.81E+07 | 2 | 4.0 |
| sp\|Q99JY0\|ECHB_MOUSE | Hadhb | Inf | 10.59 | 2.85E+06 | 0.00E+00 | 4.73E+08 | 4.47E+07 | 1 | 2.3 |
| sp\|Q62186\|SSRD_MOUSE | Ssr4 | Inf | 8.37 | 1.17E+06 | 0.00E+00 | 1.72E+09 | 2.06E+08 | 2 | 18.0 |
| sp\|O89079\|COPE_MOUSE | Cope | Inf | 5.04 | 1.42E+07 | 0.00E+00 | 1.35E+09 | 2.67E+08 | 2 | 10.7 |
| sp\|Q91W89\|MA2C1_MOUSE | Man2c1 | Inf | 4.14 | 1.36E+07 | 0.00E+00 | 1.76E+09 | 4.27E+08 | 6 | 11.8 |
| sp\|Q8VDN2\|AT1A1_MOUSE | Atp1a1 | Inf | 3.62 | 1.36E+07 | 0.00E+00 | 4.14E+09 | 1.14E+09 | 7 | 11.2 |
| sp\|P80316\|TCPE_MOUSE | Cct5 | Inf | 3.51 | 4.72E+05 | 0.00E+00 | 1.72E+09 | 4.90E+08 | 2 | 5.0 |
| sp\|Q01338\|ADA2A_MOUSE | Adra2a | Inf | 3.12 | 1.23E+06 | 0.00E+00 | 2.93E+07 | 9.38E+06 | 1 | 4.2 |
| sp\|Q9JJK2\|LANC2_MOUSE | Lancl2 | Inf | 2.96 | 3.49E+07 | 0.00E+00 | 5.71E+08 | 1.93E+08 | 3 | 10.2 |
| sp\|P51150\|RAB7A_MOUSE | Rab7a | Inf | 2.36 | 8.33E+07 | 0.00E+00 | 1.68E+09 | 7.15E+08 | 2 | 12.1 |
| sp\|Q9CZD3\|SYG_MOUSE | Gars | Inf | 2.35 | 1.71E+08 | 0.00E+00 | 2.87E+09 | 1.22E+09 | 8 | 17.7 |
| sp\|Q91VR2\|ATPG_MOUSE | Atp5c1 | Inf | 2.25 | 2.93E+08 | 0.00E+00 | 7.61E+09 | 3.38E+09 | 6 | 21.5 |
| sp\|Q61187\|TS101_MOUSE | Tsg101 | Inf | 2.21 | 5.67E+07 | 0.00E+00 | 1.59E+09 | 7.22E+08 | 2 | 7.2 |
| sp\|Q9Z1Z0\|USO1_MOUSE | Uso1 | Inf | 2.12 | 2.25E+07 | 0.00E+00 | 1.42E+09 | 6.71E+08 | 2 | 5.0 |
| sp\|O55023\|IMPA1_MOUSE | Impa1 | Inf | 2.01 | 5.55E+07 | 0.00E+00 | 2.20E+10 | 1.09E+10 | 8 | 29.6 |
| sp\|P70698\|PYRG1_MOUSE | Ctps1 | Inf | 1.94 | 4.35E+06 | 0.00E+00 | 7.37E+09 | 3.79E+09 | 10 | 25.4 |
| sp\|Q9CR00\|PSMD9_MOUSE | Psmd9 | Inf | 1.88 | 4.94E+07 | 0.00E+00 | 4.20E+08 | 2.24E+08 | 1 | 7.2 |
| sp\|P47915\|RL29_MOUSE | Rpl29 | Inf | 1.73 | 2.55E+08 | 0.00E+00 | 6.89E+08 | 3.97E+08 | 2 | 11.9 |
| sp\|Q61081\|CDC37_MOUSE | Cdc37 | Inf | 1.73 | 1.30E+07 | 0.00E+00 | 5.25E+08 | 3.04E+08 | 2 | 7.1 |
| sp\|Q91V41\|RAB14_MOUSE | Rab14 | Inf | 1.67 | 3.99E+08 | 0.00E+00 | 7.99E+09 | 4.80E+09 | 6 | 48.8 |
| sp\|Q64213\|SF01_MOUSE | Sf1 | Inf | 1.56 | 9.25E+05 | 0.00E+00 | 8.15E+08 | 5.22E+08 | 2 | 3.5 |
| sp\|Q9JLJ2\|AL9A1_MOUSE | Aldh9a1 | Inf | 1.56 | 9.29E+04 | 0.00E+00 | 6.97E+09 | 4.47E+09 | 5 | 15.6 |
| sp\|Q61753\|SERA_MOUSE | Phgdh | Inf | 1.49 | 3.29E+06 | 0.00E+00 | 2.39E+09 | 1.60E+09 | 3 | 6.8 |
| sp\|P15105\|GLNA_MOUSE | Glul | Inf | 1.40 | 1.82E+07 | 0.00E+00 | 3.81E+10 | 2.73E+10 | 16 | 50.4 |
| sp\|Q80X90\|FLNB_MOUSE | Flnb | Inf | 1.35 | 5.64E+06 | 0.00E+00 | 1.73E+09 | 1.28E+09 | 7 | 4.3 |
| sp\|Q9CVB6\|ARPC2_MOUSE | Arpc2 | Inf | 1.34 | 9.71E+06 | 0.00E+00 | 2.35E+09 | 1.75E+09 | 2 | 6.7 |
| sp\|P99029\|PRDX5_MOUSE | Prdx5 | Inf | 1.34 | 7.45E+06 | 0.00E+00 | 2.36E+08 | 1.76E+08 | 2 | 12.4 |
| sp\|P63101\|1433Z_MOUSE | Ywhaz | Inf | 1.26 | 5.21E+07 | 0.00E+00 | 1.00E+09 | 7.97E+08 | 4 | 20.4 |
| sp\|P58281\|OPA1_MOUSE | Opa1 | Inf | 1.25 | 1.89E+08 | 0.00E+00 | 2.11E+09 | 1.69E+09 | 11 | 14.5 |
| sp\|P70404\|IDHG1_MOUSE | Idh3g | Inf | 1.25 | 6.86E+06 | 0.00E+00 | 4.19E+08 | 3.35E+08 | 1 | 2.8 |
| sp\|Q9CZP0\|UFSP1_MOUSE | Ufsp1 | Inf | 1.19 | 1.15E+08 | 0.00E+00 | 4.35E+09 | 3.67E+09 | 5 | 41.5 |
| sp\|Q99LF4\|RTCB_MOUSE | Rtcb | Inf | 1.13 | 8.92E+07 | 0.00E+00 | 2.81E+09 | 2.49E+09 | 4 | 9.7 |
| sp\|Q9D1Q6\|ERP44_MOUSE | Erp44 | Inf | 1.12 | 9.57E+07 | 0.00E+00 | 1.35E+11 | 1.21E+11 | 21 | 62.3 |
| sp\|P62281\|RS11_MOUSE | Rps11 | Inf | 1.04 | 5.12E+08 | 0.00E+00 | 3.94E+09 | 3.80E+09 | 5 | 32.3 |
| sp\|Q3UTJ2\|SRBS2_MOUSE | Sorbs2 | Inf | 1.01 | 6.56E+07 | 0.00E+00 | 9.41E+08 | 9.36E+08 | 2 | 4.0 |

| Protein | Gene Name | Ratio OTUB2/Cont (FLAG) | Ratio OTUB2/Cont (GFP) | FLAGotub2 | FLAG | GFPotub2 | GFP | # peptides | Seq. Coverage |
|---|---|---|---|---|---|---|---|---|---|
| sp\|Q9JKX6\|NUDT5_MOUSE | Nudt5 | Inf | 0.99 | 6.00E+07 | 0.00E+00 | 7.65E+08 | 7.76E+08 | 1 | 6.0 |
| sp\|P28184\|MT3_MOUSE | Mt3 | Inf | 0.95 | 1.84E+07 | 0.00E+00 | 4.51E+08 | 4.75E+08 | 1 | 17.6 |
| sp\|P18872\|GNAO_MOUSE | Gnao1 | Inf | 0.90 | 3.22E+06 | 0.00E+00 | 2.42E+08 | 2.70E+08 | 1 | 4.2 |
| sp\|Q9D7G0\|PRPS1_MOUSE | Prps1 | Inf | 0.80 | 1.46E+09 | 0.00E+00 | 2.48E+09 | 3.11E+09 | 9 | 37.7 |
| sp\|Q8CG76\|ARK72_MOUSE | Akr7a2 | Inf | 0.73 | 2.58E+07 | 0.00E+00 | 8.97E+08 | 1.23E+09 | 3 | 12.8 |
| sp\|P21550\|ENOB_MOUSE | Eno3 | Inf | 0.59 | 2.97E+08 | 0.00E+00 | 9.85E+08 | 1.67E+09 | 2 | 7.6 |
| sp\|P41241\|CSK_MOUSE | Csk | Inf | 0.23 | 1.83E+07 | 0.00E+00 | 2.88E+07 | 1.26E+08 | 2 | 5.8 |
| sp\|P09602\|HMGN2_MOUSE | Hmgn2 | Inf | 0.06 | 5.63E+08 | 0.00E+00 | 1.15E+07 | 1.85E+08 | 1 | 16.7 |
| sp\|Q9DCC4\|P5CR3_MOUSE | Pycrl | Inf | 0.01 | 4.21E+08 | 0.00E+00 | 8.28E+06 | 1.04E+09 | 1 | 5.8 |
| sp\|Q8BP48\|MAP11_MOUSE | Metap1 | Inf | 0.00 | 1.03E+08 | 0.00E+00 | 0.00E+00 | 1.96E+08 | 1 | 4.9 |
| sp\|Q9CQX0\|OTUB2_MOUSE | Otub2 | 145.09 | 179.16 | 1.16E+10 | 7.97E+07 | 1.43E+11 | 7.99E+08 | 15 | 47.9 |
| sp\|P34884\|MIF_MOUSE | Mif | 99.66 | 0.98 | 2.48E+09 | 2.49E+07 | 2.12E+11 | 2.17E+11 | 6 | 51.3 |
| sp\|P06151\|LDHA_MOUSE | Ldha | 30.62 | 1.08 | 1.14E+09 | 3.72E+07 | 2.14E+10 | 1.98E+10 | 18 | 57.5 |
| sp\|P62774\|MTPN_MOUSE | Mtpn | 24.03 | 1.52 | 6.66E+08 | 2.77E+07 | 4.37E+09 | 2.88E+09 | 2 | 20.3 |
| sp\|P60843\|IF4A1_MOUSE | Eif4a1 | 22.52 | 1.10 | 9.98E+07 | 4.43E+06 | 5.19E+09 | 4.72E+09 | 7 | 22.7 |
| sp\|Q8R081\|HNRPL_MOUSE | Hnrnpl | 20.81 | 1.47 | 3.85E+07 | 1.85E+06 | 2.16E+09 | 1.47E+09 | 4 | 10.2 |
| sp\|P59999\|ARPC4_MOUSE | Arpc4 | 18.81 | 1.53 | 8.70E+06 | 4.62E+05 | 7.42E+08 | 4.85E+08 | 1 | 4.8 |
| sp\|K1C9_HUMAN\| | | 18.55 | 1.18 | 1.75E+09 | 9.42E+07 | 6.95E+09 | 5.88E+09 | 11 | 31.1 |
| sp\|Q61024\|ASNS_MOUSE | Asns | 11.89 | 1.41 | 4.05E+07 | 3.40E+06 | 2.75E+10 | 1.94E+10 | 19 | 45.6 |
| sp\|P63085\|MK01_MOUSE | Mapk1 | 11.56 | 1.26 | 1.50E+08 | 1.30E+07 | 2.15E+09 | 1.70E+09 | 4 | 12.8 |
| sp\|Q3V1L4\|5NTC_MOUSE | Nt5c2 | 10.78 | 0.71 | 1.57E+08 | 1.46E+07 | 1.78E+09 | 2.51E+09 | 6 | 12.5 |
| sp\|Q00612\|G6PD1_MOUSE | G6pdx | 8.15 | 1.47 | 1.44E+07 | 1.77E+06 | 8.79E+08 | 5.98E+08 | 2 | 6.4 |
| sp\|Q99JN2\|KLH22_MOUSE | Klhl22 | 7.85 | 0.82 | 1.62E+09 | 2.06E+08 | 2.98E+09 | 3.61E+09 | 6 | 15.8 |
| sp\|Q9D1G2\|PMVK_MOUSE | Pmvk | 7.81 | 1.42 | 5.54E+07 | 7.10E+06 | 1.62E+10 | 1.14E+10 | 10 | 38.5 |
| sp\|P32921\|SYWC_MOUSE | Wars | 7.24 | 2.08 | 2.99E+07 | 4.14E+06 | 1.48E+09 | 7.10E+08 | 2 | 5.4 |
| sp\|P97315\|CSRP1_MOUSE | Csrp1 | 7.10 | 1.32 | 4.21E+07 | 5.93E+06 | 1.44E+09 | 1.09E+09 | 1 | 8.8 |
| sp\|P62858\|RS28_MOUSE | Rps28 | 5.38 | 0.85 | 6.20E+07 | 1.15E+07 | 1.17E+10 | 1.38E+10 | 5 | 56.5 |
| sp\|Q91YQ5\|RPN1_MOUSE | Rpn1 | 5.06 | 2.45 | 9.01E+07 | 1.78E+07 | 2.11E+09 | 8.64E+08 | 3 | 5.4 |
| sp\|Q9D819\|IPYR_MOUSE | Ppa1 | 4.78 | 1.01 | 2.78E+07 | 5.82E+06 | 1.58E+08 | 1.56E+08 | 1 | 3.1 |
| sp\|P62751\|RL23A_MOUSE | Rpl23a | 4.02 | 1.20 | 9.39E+07 | 2.33E+07 | 6.86E+08 | 5.74E+08 | 2 | 15.4 |
| sp\|Q9DB29\|IAH1_MOUSE | Iah1 | 3.83 | 1.44 | 1.25E+09 | 3.26E+08 | 3.72E+10 | 2.58E+10 | 8 | 48.6 |
| sp\|Q9R0Q7\|TEBP_MOUSE | Ptges3 | 3.71 | 1.04 | 5.65E+08 | 1.52E+08 | 7.99E+09 | 7.70E+09 | 4 | 23.1 |
| sp\|Q64518\|AT2A3_MOUSE | Atp2a3 | 3.09 | 4.56 | 5.57E+07 | 1.80E+07 | 5.43E+08 | 1.19E+08 | 5 | 6.6 |
| sp\|P68040\|GBLP_MOUSE | Gnb2l1 | 2.91 | 1.57 | 2.75E+07 | 9.46E+06 | 9.87E+09 | 6.28E+09 | 12 | 46.7 |
| sp\|P84104\|SRSF3_MOUSE | Srsf3 | 2.86 | 5.26 | 1.75E+08 | 6.13E+07 | 8.31E+09 | 1.58E+09 | 2 | 12.2 |
| sp\|Q9D8E6\|RL4_MOUSE | Rpl4 | 2.58 | 1.34 | 5.31E+09 | 2.06E+09 | 1.49E+09 | 1.11E+09 | 9 | 23.4 |
| sp\|Q9DCF9\|SSRG_MOUSE | Ssr3 | 2.53 | 3.35 | 3.77E+07 | 1.49E+07 | 1.49E+09 | 4.45E+08 | 1 | 7.6 |
| sp\|P14148\|RL7_MOUSE | Rpl7 | 2.44 | 1.36 | 7.25E+08 | 2.97E+08 | 4.18E+09 | 3.07E+09 | 7 | 20.7 |
| sp\|Q6PHN9\|RAB35_MOUSE | Rab35 | 2.37 | 1.62 | 6.88E+08 | 2.90E+08 | 4.45E+09 | 2.75E+09 | 1 | 5.5 |
| sp\|P62983\|RS27A_MOUSE | Rps27a | 2.34 | 0.90 | 1.45E+08 | 6.22E+07 | 2.27E+06 | 2.51E+06 | 2 | 14.1 |
| sp\|P62918\|RL8_MOUSE | Rpl8 | 2.32 | 2.04 | 1.99E+09 | 8.57E+08 | 2.35E+09 | 1.15E+09 | 5 | 18.3 |

| Protein | Gene Name | Ratio OTUB2/Cont (FLAG) | Ratio OTUB2/Cont (GFP) | FLAGotub2 | FLAG | GFPotub2 | GFP | # peptides | Seq. Coverage |
|---|---|---|---|---|---|---|---|---|---|
| sp\|P63017\|HSP7C_MOUSE | Hspa8 | 2.14 | 1.60 | 2.23E+09 | 1.04E+09 | 1.97E+10 | 1.23E+10 | 22 | 35.6 |
| sp\|P62717\|RL18A_MOUSE | Rpl18a | 2.11 | 1.17 | 1.38E+09 | 6.54E+08 | 7.78E+08 | 6.67E+08 | 2 | 10.8 |
| sp\|P47962\|RL5_MOUSE | Rpl5 | 2.02 | 2.57 | 4.24E+08 | 2.10E+08 | 9.92E+08 | 3.86E+08 | 3 | 9.4 |
| sp\|Q68FD5\|CLH1_MOUSE | Cltc | 1.84 | 1.53 | 2.55E+08 | 1.38E+08 | 5.10E+09 | 3.32E+09 | 11 | 9.6 |
| sp\|O88533\|DDC_MOUSE | Ddc | 1.76 | 1.62 | 1.98E+08 | 1.13E+08 | 1.47E+10 | 9.07E+09 | 11 | 29.8 |
| sp\|Q60864\|STIP1_MOUSE | Stip1 | 1.73 | 0.48 | 9.35E+07 | 5.42E+07 | 2.42E+08 | 5.08E+08 | 2 | 4.2 |
| sp\|Q91VR5\|DDX1_MOUSE | Ddx1 | 1.71 | 1.99 | 1.09E+08 | 6.38E+07 | 1.07E+09 | 5.38E+08 | 2 | 5.3 |
| sp\|Q9JMD3\|PCTL_MOUSE | Stard10 | 1.64 | 0.00 | 4.18E+07 | 2.55E+07 | 0.00E+00 | 3.14E+08 | 1 | 4.5 |
| sp\|P62960\|YBOX1_MOUSE | Ybx1 | 1.56 | 0.00 | 1.51E+08 | 9.72E+07 | 0.00E+00 | 7.84E+07 | 1 | 7.5 |
| sp\|P46425\|GSTP2_MOUSE | Gstp2 | 1.51 | 0.62 | 1.08E+10 | 7.20E+09 | 4.60E+09 | 7.46E+09 | 17 | 74.3 |
| sp\|P62830\|RL23_MOUSE | Rpl23 | 1.51 | 0.00 | 2.10E+08 | 1.39E+08 | 7.88E+05 | 2.84E+08 | 2 | 20.0 |
| sp\|Q9JJV2\|PROF2_MOUSE | Pfn2 | 1.49 | 6.72 | 2.41E+07 | 1.62E+07 | 1.19E+10 | 1.76E+09 | 7 | 46.4 |
